# Pathogen-induced formation of a nascent organelle derived from mitochondria

**DOI:** 10.64898/2026.04.23.720395

**Authors:** Xianhe Li, José M. Delgado, Yubai Sun, Lena Pernas

**Author notes:** Department of Microbiology & Infectious Disease Center, School of Basic Medical Sciences, Peking University Health Science Center, Beijing, China.

## Abstract

Intracellular pathogens extensively remodel host cells to create environments that support their survival and replication. How pathogens manipulate host organelle biology to promote infection remains a central question in microbial pathogenesis. Here we show that the human parasite *Toxoplasma gondii* induces the biogenesis of acidified compartments derived from host mitochondria that promote parasite growth. Following infection, host mitochondria shed large <u>s</u>tructures <u>p</u>ositive for <u>o</u>uter mitochondrial membrane (OMM), termed SPOTs. SPOTs matured into multivesicular compartments that engulfed cytosolic protein and functional lysosomes. The acquisition of host lysosomes by SPOTs required host ESCRT machinery and the parasite effector TgGRA7, and drove the acidification of the SPOT lumen. Disrupting SPOT acidification impaired parasite proliferation, implicating SPOT maturation in parasite fitness. These findings show that an intracellular pathogen co-opts host machinery to drive the formation of a nascent organelle derived from mitochondria, and raise the possibility that mitochondria can be reprogrammed to generate new organelles with specialized functions.

**One-Sentence Summary:** An intracellular pathogen hijacks host mitochondrial membranes and ESCRT machinery to generate an acidified organelle that supports parasite replication.

## Main Text

To establish nutrient-rich replicative niches and evade innate immune defenses, intracellular pathogens use diverse strategies to remodel cellular membranes, including through the reprogramming of membrane trafficking and the hijacking of the endolysosomal system ^1,2^. Although how pathogens manipulate preexisting host endolysosomal membranes is well characterized, whether and how pathogens exploit host mitochondrial membranes remains poorly understood.

Mitochondria possess a uniquely complex ultrastructure, with an inner mitochondrial membrane and matrix encased by a single outer mitochondrial membrane (OMM) ^3^. The OMM has long been considered a relatively passive barrier and platform for solute exchange, in contrast to the intricate inner membranes that play a key role in regulating mitochondrial metabolic function ^4^. However, this static view of the OMM has been challenged by recent studies showing that it is actively remodeled to sustain peroxisome biogenesis, and gives rise to membrane-bound structures during cellular stress and infection with the human parasite *Toxoplasma gondii* ^5–9^.

Here, we show that infection-induced <u>s</u>tructures <u>p</u>ositive for <u>o</u>uter mitochondrial membrane, termed SPOTs, developed into organelle-like compartments that diverged markedly from their mitochondrial progenitors in both ultrastructure and function. Over the course of infection, SPOTs matured into multivesicular compartments that internalized cytosolic protein and engulfed functional lysosomes. This maturation required host ESCRT machinery and the secreted *Toxoplasma* effector TgGRA7, and culminated in the acidification of the SPOT lumen. Impairing SPOT acidification led to reduced parasite growth, suggesting that SPOT maturation promotes *Toxoplasma* replication. Together, these findings show that an intracellular pathogen exploits the plasticity of the OMM to generate an organelle-like compartment with distinct properties—and thus that mitochondrial membranes can be repurposed to create new cellular compartments.

### SPOTs acquire emergent properties following release from mitochondria

Following invasion with *Toxoplasma gondii,* host mitochondria become tethered to the parasite vacuole (PV) through the binding of the parasite effector TgMAF1 to the host import receptor TOM70 and shed SPOTs as early as 6 hours post infection (hpi) ^10–12^. To determine whether SPOTs—like other previously reported OMM-derived structures, mitochondrial-derived vesicles (MDVs) and mitochondrial-derived compartments (MDCs)—are delivered to degradative intermediates, we first generated SPOT reporter HeLa cells expressing enhanced green fluorescent protein (eGFP) fused to the OMM-targeting transmembrane (TM) domain of OMP25 (OMM-GFP), and blue FP (BFP) fused to a matrix-targeting sequence ^5–9^. Because SPOTs retain OMM proteins but exclude matrix and inner mitochondrial membrane (IMM), this system allowed us to distinguish SPOTs from mitochondria. Next, we infected our SPOT reporter cells with mCherry-expressing *Toxoplasma* parasites (*Toxo^mCh^*) and asked if SPOTs were trafficked to autophagosomes and multivesicular bodies (MVBs), organelles that converge on the lysosome for degradation. We found that SPOTs lacked LC3 and CD63, markers of autophagosomes and MVBs, respectively (**Extended Data Fig. 1**). Rather, over the course of infection SPOTs increased in number, diameter, and became multivesicular (**Fig. 1a-g**). By 24 hpi, over 40% of SPOTs contained 2 or more internal vesicles (**Fig. 1g**). No SPOTs were observed in uninfected cells at the corresponding time points (**Fig. 1d**). Thus, SPOTs are not delivered to endolysosomal trafficking intermediates but instead develop into larger and multivesicular structures during infection ^10^.

**Figure 1:**
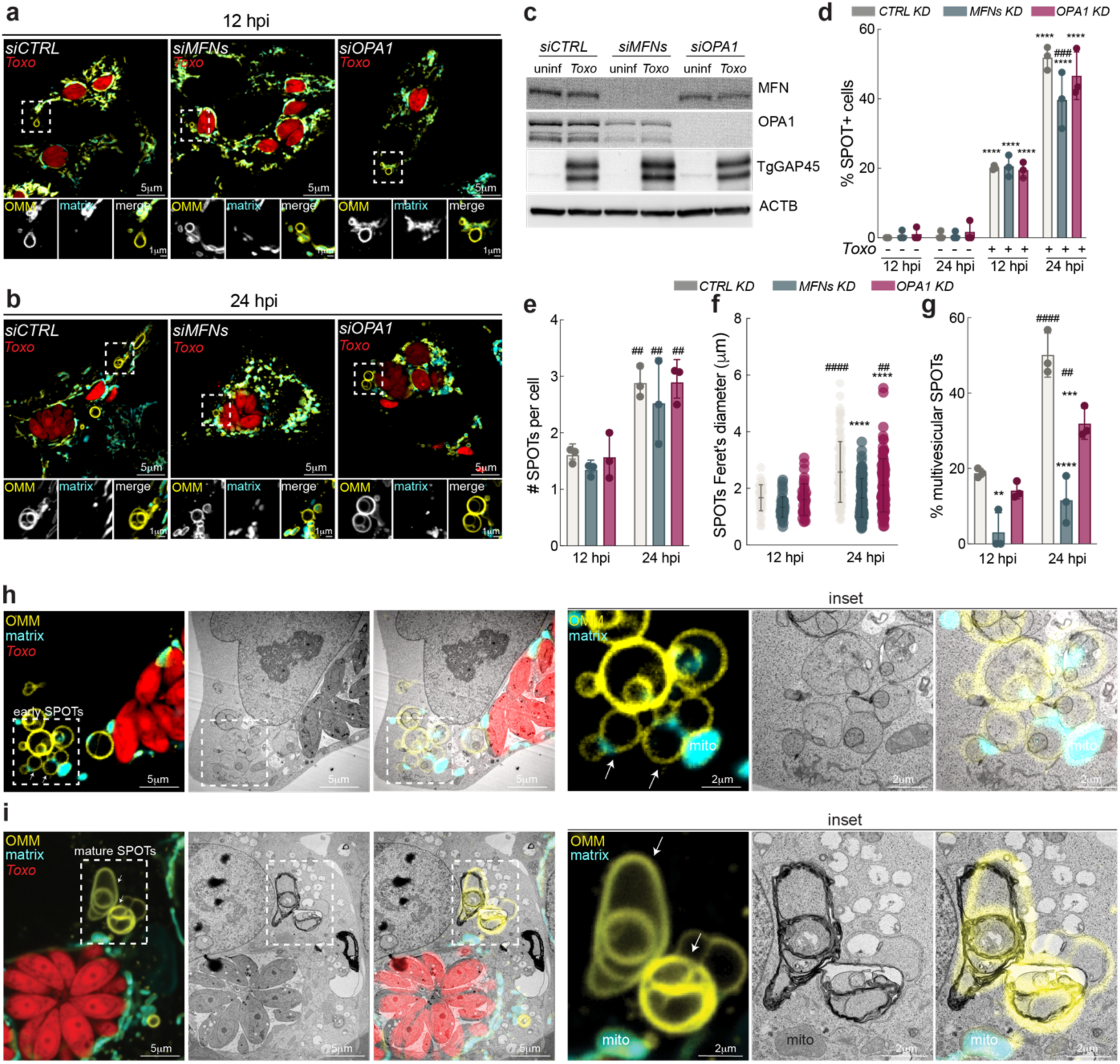
SPOTs develop into compartmentalized structures. Representative live-cell images of *CTRL, MFN1&2,* and *OPA1-*silenced (si) HeLa cells expressing OMM-targeted GFP and matrix-targeted BFP (SPOT reporter) infected with *Toxoplasma-*mCherry (*Toxo^mCh^*) at (**a**) 12 hours post infection (hpi) and (**b**) 24 hpi; scale bar 5μm, inset 1μm. (**c**) WT, *MFN1 & 2 (MFNs)* and *OPA1*-silenced SPOT reporter HeLa cells as in (a-b) were analyzed by means of immunoblotting for MFN ∼80kDa; and ∼OPA1, 80-100 kDa; TgGAP45 ∼45 kDa. (**d**) Percentage (%) of SPOT-positive cells, (**e**) number (#) of SPOTs per cell, (**f**) SPOT diameter, and (**g**) % of multivesicular SPOTs from experiments as in (a-b); data are mean ± SEM from more than 30 infected cells from three replicates; ## p<0.01, ### p<0.001, #### p<0.0001 for 12 hpi versus 24 hpi, **p<0.01, ***p<0.001, ****p<0.0001 for *siCTRL* versus *siMFN, siOPA1* by means of two-way ANOVA analysis. Correlative light electron microscopy (CLEM) analysis of *Toxo^mCh^*- infected SPOT reporter cells with (**h**) early SPOTs and (**i**) mature SPOTs indicated by white arrows; scale bar 5μm, inset 2μm.

How do SPOTs grow and develop structural complexity? During infection, the OMM proteins mitofusin 1 (MFN1) and MFN2, which tether mitochondria to each other to mediate fusion, are redistributed to SPOT membranes ^3,10^. We therefore reasoned that MFNs, as for mitochondria, regulate SPOT dynamics. However, the possibility remained that SPOT growth simply reflected the release of larger SPOTs at late stages of infection. To distinguish between these possibilities, we compared SPOT morphology at 12 and 24 hpi in *Toxo^mCh^*-infected SPOT reporter cells in which *MFN1* and *MFN2* were silenced using small interfering RNAs (siRNAs). The siRNA depletion of MFN1 and MFN2 significantly blocked the growth in SPOT diameter from 12 hpi to 24 hpi and prevented the formation of multivesicular SPOTs (**Fig. 1a-g**). *MFN1* and *MFN2* ablation did not, however, affect SPOT frequency or number during infection (**Fig. 1a-e**). To distinguish whether MFNs promoted SPOT growth through membrane fusion or through a fusion-independent mechanism, we also examined SPOTs in cells lacking the IMM protein OPA1. As in MFN-deficiency, cells lacking OPA1 exhibit small, fragmented mitochondria due to impaired fusion (**Fig. 1a-b**) ^3^. In contrast to MFN-deficient cells, SPOTs in cells treated with *siOPA1* significantly increased in size during infection and became multivesicular, albeit to a lesser extent than in *siCTRL-*treated cells (**Fig. 1a-g**). Thus, SPOTs require MFN1 and MFN2 to increase in size and become multivesicular in a fusion-independent manner, consistent with a role for MFNs in SPOT tethering rather than membrane fusion.

### SPOTs are compartmentalized and contain diverse cargo

We next asked whether the progression of SPOTs into large, multivesicular structures—which we will hereinafter refer to as maturation—was concomitant with changes in their content. To address this question, we turned to correlative light electron microscopy (CLEM), which enabled a comparison of the ultrastructure and contents of early SPOTs to mature SPOTs in an unbiased manner. We confirmed that mature SPOTs were multilamellar and multivesicular, in contrast to early SPOTs (**Fig. 1h-i**). Our CLEM analysis also revealed two unexpected features of SPOTs. First, we found that both early and mature SPOT content resembled that of the cytosol (**Fig. 1h-i**). Second, that the cargo of vesicles within mature SPOTs was heterogeneous and differed from that of early SPOTs (**Fig. 1h-i**). Thus, SPOT maturation is accompanied by changes in both morphology and cargo.

Surprisingly, we also found that membrane-bound structures reminiscent of lysosomes were readily observed within intralumenal vesicles of mature SPOTs (**Fig. 1i**). To confirm that the structures within SPOTs were indeed lysosomes, we used CLEM to analyze SPOT reporter cells infected with *Toxo^mCh^* parasites and labeled with LysoTracker, a dye that accumulates in lysosomes in a pH-dependent manner and thus labels acidified lysosomes. Examination of a mature SPOT revealed three lysosomes and other membrane-bound structures of unknown origin (**Fig. 2a**). In addition to retaining an acidic pH, lysosomes within SPOTs were proteolytically active as assessed by the Magic Red substrate, which fluoresces upon cleavage by the lysosomal protease cathepsin B (**Extended Data Fig. 2**). Using the lysosomal membrane protein LAMP2, we found that more than 40% of SPOTs at 24 hpi contained lysosomes, in contrast to less than 10% of SPOTs at 12 hpi (**Fig. 2b-c**). Thus, SPOT maturation is associated with the accumulation of proteolytically active lysosomes during infection.

**Figure 2:**
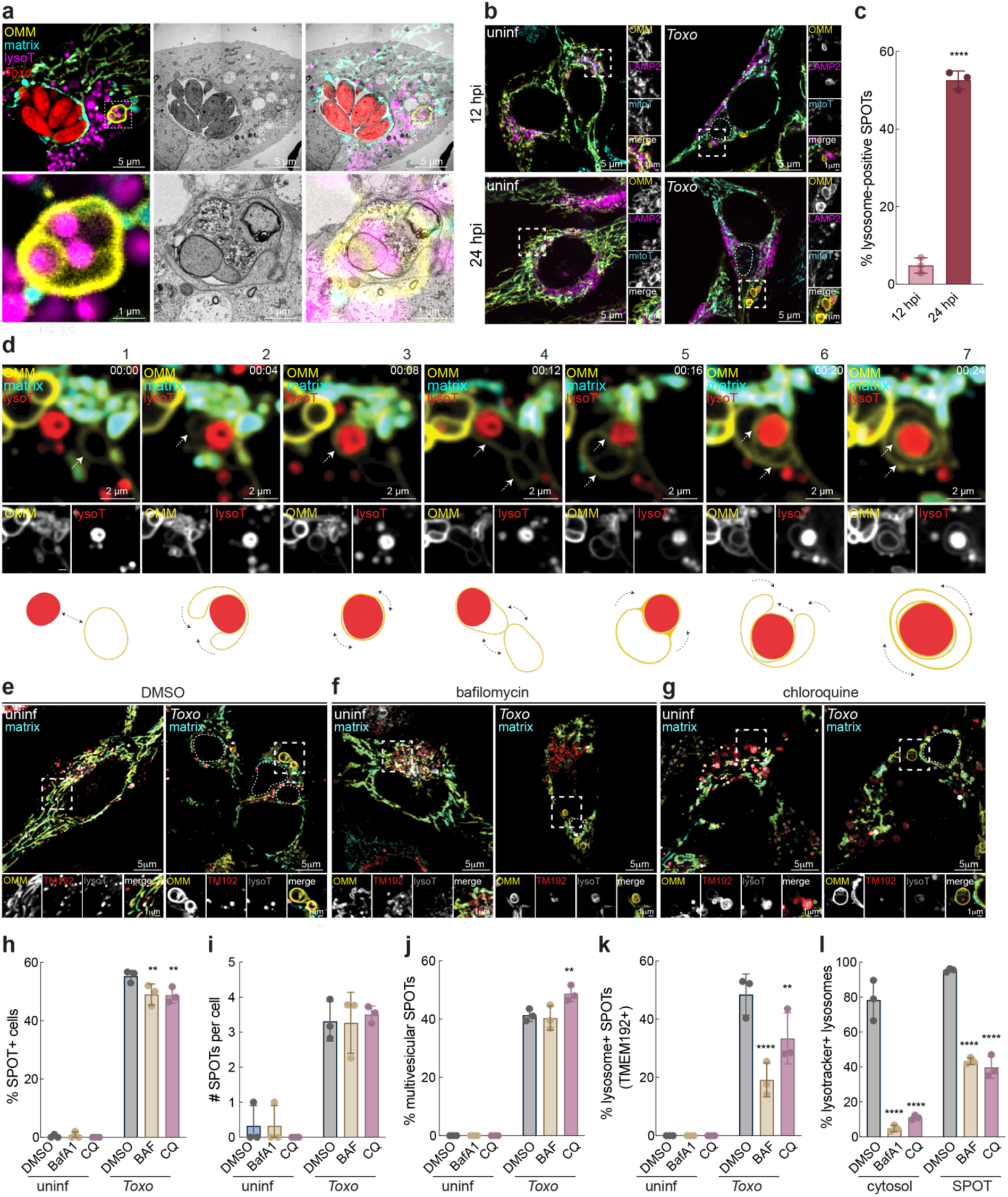
SPOTs engulf active lysosomes. (**a**) CLEM analysis of *Toxo^mCh^*-infected SPOT reporter cells labeled with lysotracker (lysoT); Inset of a SPOT positive for three lysosomes; scale bars 5 μm and (inset) 1 μm. (**b**) Representative immunofluorescence (IF) images of uninf and *Toxo*-infected SPOT reporter Hela cells at 12 hpi and 24 hpi. (inset) SPOTs in *Toxo-*infected cells contain a lysosome (LAMP2) at 24 hpi, but not 12 hpi; scale bars 5 μm and (inset) 1 μm. (**c**) Percentage (%) of SPOTs containing lysosomes from images as in (B), data are mean ± SEM from more than 30 infected cells from three replicates, ****p<0.0001 by unpaired t-test. (**d**) Live cell images of SPOT reporter cells infected with *Toxoplasma* and labeled with lysotracker at 24 hpi; white arrows indicate a SPOT that engulfs a lysosome at the indicated time frames with a schematic representation below. (**e**) Representative live-cell images of uninfected (uninf) and *Toxo*-infected SPOT reporter cells that were treated with DMSO; (**f**) bafilomycin; and (**g**) chloroquine at 18 hpi and imaged at 24 hpi; scale bars 5 μm, inset 2 μm. TMEM192 (TM192) (**h**) Percentage (%) of SPOT-positive cells; (**i**) number of SPOTs per cell; (**j**) % of multivesicular SPOTs; (**k**) % of lysosome-containing SPOTs, and (**l**) % of lysotracker-positive lysosomes in cytosol or SPOTs from experiments as in (e-g); data are mean ± of 30 or more infected cells from three replicates; **p<0.01; ****p<0.0001 for vehicle versus treatment by means of two-way ANOVA analysis.

### SPOTs engulf functional lysosomes during infection

How do OMM-derived structures acquire lysosomes, rather than being degraded by them as occurs in mitophagy? We reasoned that SPOTs may acquire lysosomes through either fusion, as occurs between lysosomes and autophagosomes during autophagy, or engulfment. To distinguish between these scenarios, we silenced *STX17* and *RAB7*, key mediators of lysosome-autophagosome fusion, in *Toxo^mCh^-*infected SPOT reporter cells ^13,14^. However, neither the loss of STX17 nor RAB7 inhibited SPOT uptake of lysosomes (**Extended Data Fig. 3**). Consistent with a fusion-independent process, lysosomal membrane proteins LAMP2 and the vacuolar type ATPase (v-ATPase) subunit ATP6V0D1 were excluded from membranes of SPOTs containing lysosomes (**Fig. 2b; Extended Data Fig. 4a**). Furthermore, SPOTs contained endogenous levels of GAPDH and cytosolic GFP, neither of which would be expected if SPOTs fused with lysosomes (**Extended Data Fig. 4b-i**). To capture the acquisition process, we performed live-cell imaging of infected SPOT reporter cells at 24 hpi, when lysosome-positive SPOTs were most frequently observed (**Fig. 2b-c**). Over a 24-minute time course, we observed SPOT membranes dynamically extending around and engulfing an acidified lysosome (**Fig. 2d**). This led to the enclosure of the lysosome within multiple layers of SPOT membrane, consistent with the concentric membranes observed in our CLEM analysis (**Fig. 1h**; **Fig. 2d**). Thus, SPOT maturation is concomitant with the engulfment of acidified lysosomes.

We noted that lysosomes were four times more likely to be found within SPOTs than similarly sized organelles including peroxisomes and early endosomes (**Extended Data Fig. 5**). Because a unique feature of lysosomes is their low pH, we asked whether lysosomal acidification was required for SPOT uptake of lysosomes. To do so, we turned to the lysosomotropic agents bafilomycin-A1 (BafA1) and chloroquine (CQ), which inhibit v-ATPase—a molecular pump that uses ATP hydrolysis to drive protons against their electrochemical gradient into the lumen of the lysosome—and thus impair lysosomal acidification ^15,16^. To visualize lysosomes independently of pH, we generated SPOT reporter cells expressing the lysosomal membrane protein TMEM192 fused to RFP ^17^. Following infection, these cells were incubated with either vehicle, BafA1 or CQ (**Fig. 2e-g**). BafA1 and CQ treatment led to a significant decrease in SPOTs positive for lysosomes, despite SPOTs forming at a similar frequency (**Fig. 2e-k**). Consistent with a role for pH in mediating lysosome uptake by SPOTs, we found that, although BafA1- and CQ-treatment reduced the percentage of acidified lysosomes in treated cells to <10%, ∼40% of lysosomes within SPOTs were acidified (**Fig. 2e-g, 2l**). Thus, lysosomal acidification is required for their uptake by SPOTs.

### ESCRT machinery is enriched in isolates of mitochondria tethered to the *Toxoplasma* vacuole

How do SPOTs become multivesicular and engulf lysosomes even though their mitochondrial precursors do not? Because SPOTs emerge from mitochondria that are tethered to the *Toxoplasma* vacuole, we reasoned that factors present at the interface between mitochondria and the parasite vacuole (PV) influenced SPOT maturation ^10^. To address this possibility, we leveraged the fact that the mitochondria that release SPOTs are stably tethered to the *Toxoplasma* PV via the effector TgMAF1 and the host OMM import receptor TOM70, and thus that TgMAF1 could serve as a handle to selectively enrich PV-tethered mitochondria ^10,12,18^. To this end, we infected HeLa cells expressing an OMM-targeted myc-tag (myc-OMM) with *Δmaf1* parasites engineered to express HA-tagged TgMAF1 (HA-TgMAF1). HA-TgMAF1-bound mitochondria (*Toxo-*mito) were first isolated using an HA-based magnetic bead system ^19^. The remaining mitochondria (cyto-mito) were similarly purified using the myc-OMM tag (**Fig. 3a**) ^19^. Immunoblot analysis confirmed that TgMAF1 and TOM70 were enriched in the *Toxo*-mito fraction but depleted from cyto-mito, as expected (**Fig. 3b**). In contrast, other markers of the OMM, IMM, or matrix were similarly abundant between *Toxo-*mito and cyto-mito (**Fig. 3b**). Thus, this approach enables the isolation of SPOT-generating mitochondria and associated PV membranes.

**Figure 3:**
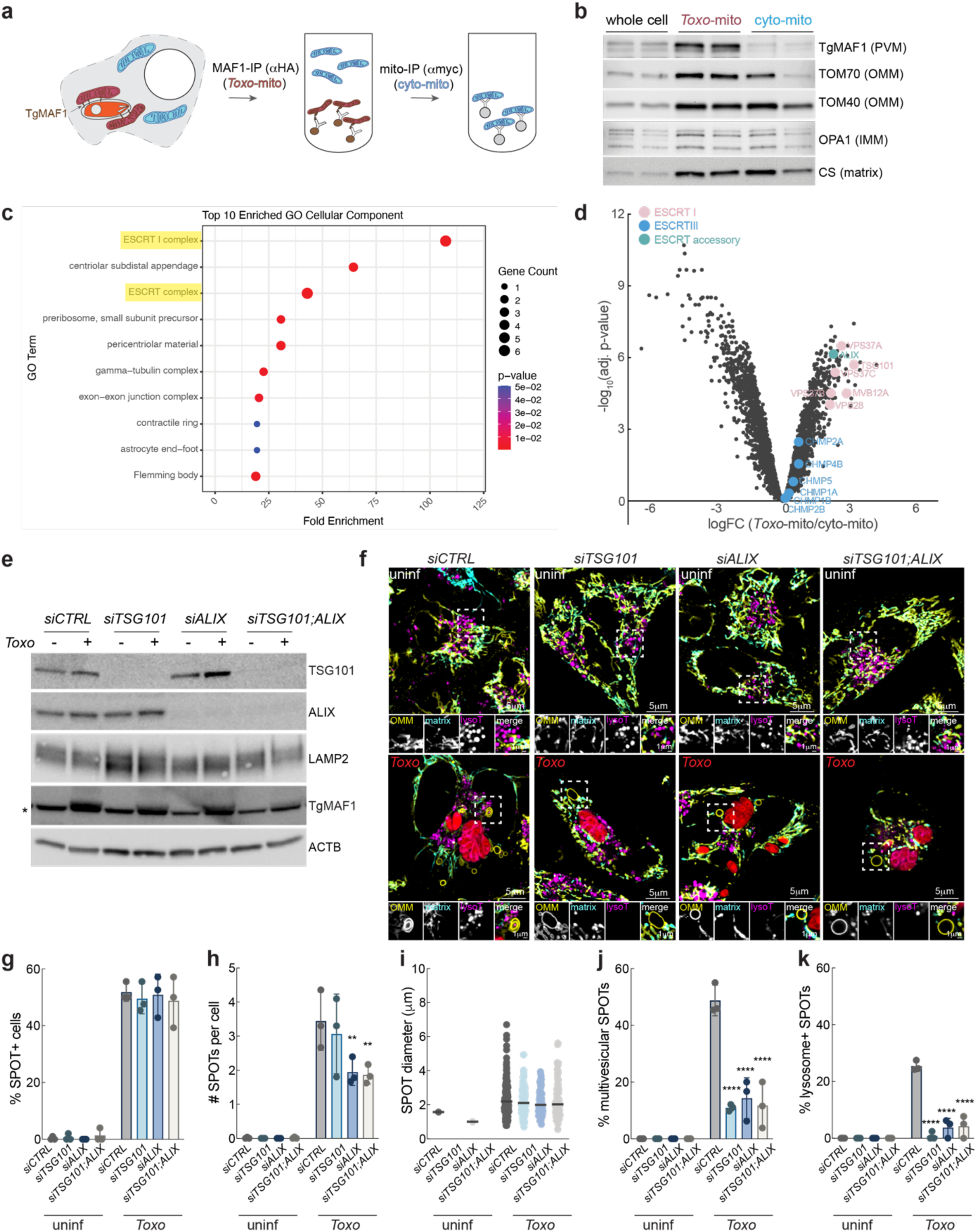
Host ESCRT machinery mediates SPOT engulfment of lysosomes. (**a**) Schematic of workflow to immunopurify PVM-associated mitochondria (*Toxo*-mito, maroon) from unassociated mitochondria (cyto-mito, blue). (**b**) *Toxo*-mito and cyto-mito were immunopurified from 3Xmyc-eGFP-OMP25-expressing HeLa cells that were infected with *Δmaf1:HA-MAF1 Toxoplasma.* Whole cell, *Toxo*-mito, and cyto-mito fractions were analyzed by means of immunoblotting for TOM70 ∼70kDa; TOM40 ∼40kDa; OPA1 80kDa; TgMAF1 ∼55 kDa; and CS ∼45 kDa. (**c**) GO Term analysis was performed using GO pathway enrichment analyses on the top 100 differentially abundant proteins between *Toxo-*mito and cyto-mito from experiments performed as in (a-b) and analyzed by mass spectrometry; *Toxo*-mito (n=5) and cyto-mito (n=4) (**d**) Volcano plot of proteins identified in *Toxo*-mito and cyto-mito fractions. ESCRT I proteins (pink); ESCRT III proteins (blue); and ESCRT accessory proteins (teal) enriched in *Toxo*-mito fraction are highlighted. (**e**) *CTRL, TSG101, ALIX, and TSG101 & ALIX-*silenced (si) SPOT reporter HeLa cells as were analyzed by means of immunoblotting for TSG101 ∼45kDa; ALIX ∼90kDa; LAMP2 ∼100kDa; TgMAF1 ∼55kDa (asterisk indicates nonspecific band); and ACTB ∼45 kDa. (**f**) Representative live-cell images of *CTRL, TSG101, ALIX, and TSG101 & ALIX-*silenced HeLa SPOT reporter cells infected with *Toxoplasma-*mCherry (*Toxo^mCh^*) and labeled with lysotracker (lysoT) at 24hpi; scale bar 5 μm, inset 1μm. (**g**) Percentage (%) of SPOT-positive cells, (**h**) number (#) SPOTs per cell, (**i**) SPOT diameter, (**j**) % of multivesicular SPOTs, and (**k**) % of lysoT-positive SPOTs from experiments as in (E-F); data are mean ± SEM from more than 30 infected cells from three replicates; **p<0.01; ****p<0.0001 for *CTRL* versus silenced by means of two-way ANOVA analysis.

We next used mass spectrometry to compare the protein abundances between *Toxo*-mito and cyto-mito immunopurified from infected cells, as well as mitochondria immunopurified from uninfected cells (uninf-mito). Among the most differentially enriched proteins in the *Toxo*-mito fraction relative to the cyto-mito or uninf-mito, gene ontology (GO) analysis revealed significant enrichment of ESCRT-I complex components (**Fig. 3c-d**, **Table 1, Extended Data Fig. 6**). At least one member of each core ESCRT-I subunit was increased in abundance in the *Toxo-*mito fraction, including TSG101 (VPS23), VPS28, VPS37A,B,C,D and MVB12A (**Fig. 3d**, **Table 1**)^20^. The ESCRT-associated adaptor ALIX was also present at approximately fourfold higher levels in the *Toxo*-mito fraction, consistent with its previous identification at the PVM (**Fig. 3d**, **Table 1**) ^21^. Thus, the ESCRT-I complex is enriched at the interface between the PVM and the mitochondria that generate SPOTs.

### Host ESCRT machinery drives SPOT maturation

The enrichment of ESCRT-I components in the *Toxo*-mito fraction was interesting to us because a unique function of ESCRT machinery is to carry out reverse-topology membrane scission through a reaction that begins with the recruitment of ESCRT-I and/or the accessory protein ALIX^20^. This function is most well-characterized during intralumenal vesicle budding into endosomes and the subsequent biogenesis of MVBs ^20,22^. Analogous to MVBs, intralumenal vesicles were observed accumulating within a SPOT that we tracked using OMM-targeted photoactivatable GFP (**Extended Data Fig. 7**). These results implicated a role for ESCRT machinery in SPOT development during infection.

To test this possibility, we infected cells depleted of either the core ESCRT-I subunit TSG101 or ALIX, both of which play central roles in recruiting and assembling downstream ESCRT machinery ^20^. We found that the formation of multivesicular SPOTs and their acquisition of lysosomes was impaired by the loss of either TSG101 or ALIX, and was not further exacerbated by their simultaneous depletion, suggesting they act in the same pathway to drive SPOT maturation (**Fig. 3g-k**). Notably, SPOT formation and overall size were comparable between WT and *TSG101-* and *ALIX*-depleted cells, indicating that ESCRT function is required for SPOT maturation but not growth (**Fig. 3g-i**). In MFN1- and MFN2-deficient cells, however, SPOTs were significantly smaller and had significantly fewer lysosomes, while the loss of OPA1 produced intermediate-sized SPOTs with modest defects in lysosome uptake (**Extended Data Fig. 8**). These findings suggest that SPOTs must reach a critical size to enable lysosome acquisition, after which ESCRT-dependent remodeling drives their maturation into multivesicular, lysosome-containing structures.

### Mature SPOTs become acidified

How does lysosome acquisition affect the biochemical properties of SPOTs? Because a key feature of a subset of mature SPOTs was the accumulation of active lysosomes, we asked whether these SPOTs developed into acidified compartments at late stages of infection. To test this possibility, we needed to monitor SPOTs independently of eGFP, which is quenched in the low pH environment of the lysosome ^23^. We therefore used a tandem fluorescence (tf) reporter comprising RFP, which retains fluorescence in a low pH environment, and eGFP fused to the transmembrane domain of Fis1 for OMM targeting (OMM-tf) in cells expressing matrix-targeted BFP (**Fig. 4a**) ^23,24^. Following infection, we found that SPOTs with both RFP and GFP fluorescence lacked lysosomes (**Extended Data Fig. 9a-b**). By contrast, structures with decreased GFP relative to RFP fluorescence were positive for the lysosomal marker LAMP2 (**Extended Data Fig. 9a-b**). We therefore classified these structures as acidified SPOTs.

**Figure 4:**
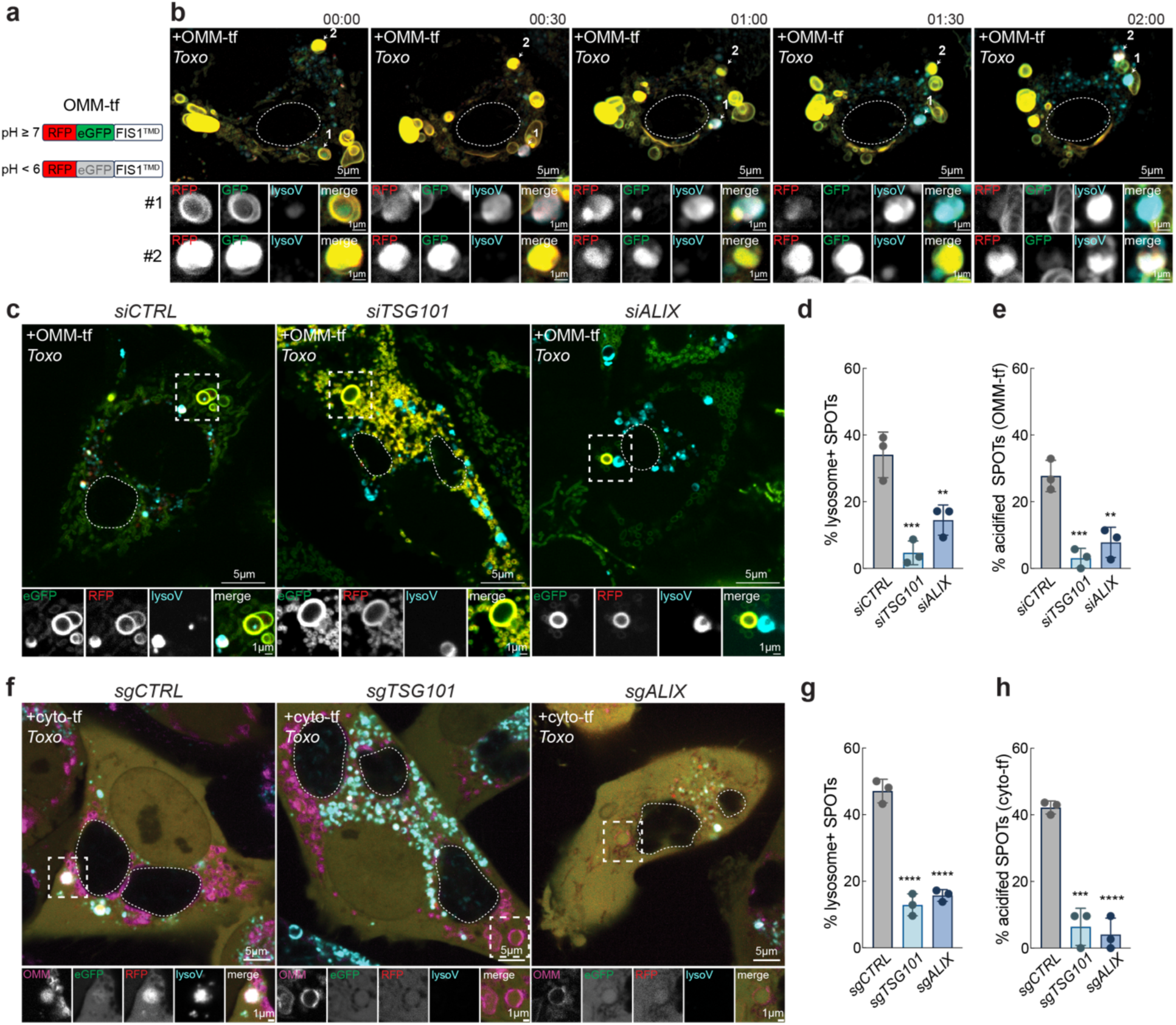
Host ESCRT machinery mediates SPOT engulfment of lysosomes. (**a**) Schematic of the OMM tandem fluorescent reporter (OMM-tf): RFP and eGFP are targeted to the OMM by the transmembrane domain of Fis1; at pH≥7 both RFP and eGFP fluoresce, at pH<6, GFP is quenched. (**b**) Live cell images of HeLa cells stably expressing OMM-tf infected with *Toxoplasma* and labeled with lysoview (lysoV) at 24 hpi; white arrows indicate two SPOTs (#1, #2) that become acidified following engulfment of a lysosome at indicated time frames post-imaging (hh:mm); dotted circle denotes parasite vacuole. (**c**) Representative live-cell images of *CTRL, TSG101, ALIX-*silenced HeLa cells expressing OMM-tf infected with *Toxoplasma* (indicated with dotted lines) and labeled with lysoview at 24 hpi; scale bars 5 μm and (inset) 1 μm. (**d**) Percentage (%) of lysoview-positive SPOTs and (**e**) % of acidified SPOTs based on OMM-tf reporter (RFP+, GFP-) from experiments as in (c); data are mean ± SEM from more than 30 infected cells from three replicates; ***p<0.001; ****p<0.0001 for *CTRL* versus silenced by means of one-way ANOVA analysis. (**f**) Representative live-cell images of *CTRL, TSG101, ALIX-*silenced HeLa cells expressing a cytosolic RFP-GFP (cyto-tf) infected with *Toxoplasma* (indicated with dotted lines) and labeled with lysoview at 24 hpi; scale bars 5μm and (inset) 1μm. (**g**) Percentage (%) of lysoview-positive SPOTs and (**h**) % of acidified SPOTs based on cyto-tf reporter (RFP+, GFP-) from experiments as in (f); data are mean ± SEM from more than 30 infected cells from three replicates; ***p<0.001; ****p<0.0001 for *CTRL* versus silenced by means of one-way ANOVA analysis.

To test whether lysosome acquisition preceded SPOT acidification, we used live-cell imaging to track two SPOTs with similar RFP and GFP levels, only one of which contained a lysosome (**Fig. 4b**). Within 30 minutes, GFP signal in the lysosome-positive SPOT was quenched (**Fig. 4b**). The SPOT initially lacking a lysosome retained GFP fluorescence, but became acidified only upon subsequent lysosome acquisition, further supporting a causal relationship between the two events (**Fig. 4b**). We next assessed the abundance of acidified SPOTs in *TSG101*- or *ALIX*- silenced cells in which lysosome uptake by SPOTs was impaired (**Fig. 3e-k**). The loss of TSG101 or ALIX reduced the frequency of acidified SPOTs by approximately 88% and 70%, respectively (**Fig. 4c-e**). Thus, lysosome uptake is required for SPOT acidification.

This finding prompted us to ask whether SPOTs acquire defining features of a lysosome, such as the capacity to acidify luminal content. To address this question, we leveraged our previous finding that SPOTs contained cytosolic protein (**Fig. 1h-i; Extended Data Fig. 4**). We generated cells co-expressing a cytosolic tandem fluorescence reporter (cyto-tf) analogous to the OMM-tf reporter, and OMM-targeted BFP to simultaneously visualize SPOT membranes ^23,24^. At 24 hpi, more than 40% of SPOTs lacked GFP but retained RFP fluorescence, indicating acidification of acquired cytosolic protein (**Fig. 4f-h**; **Extended Data Fig. 9c-d)**. By contrast, SPOTs in TSG101- or ALIX-deficient cells retained both GFP and RFP fluorescence (**Fig. 4f-h**). Thus, SPOTs acidify acquired cytosolic protein.

### Lysosome acquisition by SPOTs requires a secreted parasite factor

We reasoned that defining how ESCRT is recruited to SPOTs could provide insight into whether SPOT acidification reflected a host- or parasite-driven remodeling process. The loss of HRS, a component of ESCRT-0 that acts upstream of ESCRT-I and/or ALIX, reduced SPOT maturation to a similar extent as TSG101 or ALIX deficiency (**Extended Data Fig. 10**) ^25^. However, pharmacological inhibition of phosphatidylinositol 3-phosphate (PI3P) synthesis or ubiquitylation did not block SPOT maturation (**Extended Data Fig. 11-12**) ^20^.

Having excluded canonical host signals that recruit ESCRT-0, we next considered whether parasite-derived factors contributed to SPOT maturation ^26^. We first focused on the secreted effector TgGRA14, which promotes TSG101 recruitment proximal to the *Toxoplasma* PV ^27^. However, no differences were detected between SPOT formation, size, or uptake of lysosomes in cells infected with WT and *Δgra14* parasites (**Extended Data Fig. 13**). Given that the *Toxoplasma* effector TgMAF1 drives the formation of SPOTs, we asked whether TgMAF1-interacting proteins mediated their maturation. To this end, we examined TgMAF1 immunopurifications (IPs) from cells infected with *Δmaf1* parasites expressing hemagglutinin (HA)-tagged TgMAF1 (*Δmaf1*:HA-MAF1), which were previously analyzed by mass spectrometry ^26^. Among the most abundant parasite proteins identified were TgMAG1 and TgGRA7, effectors secreted from dense granule organelles with TgMAF1 (**Fig. 5a**; **Table 2**) ^28,29^.

**Figure 5:**
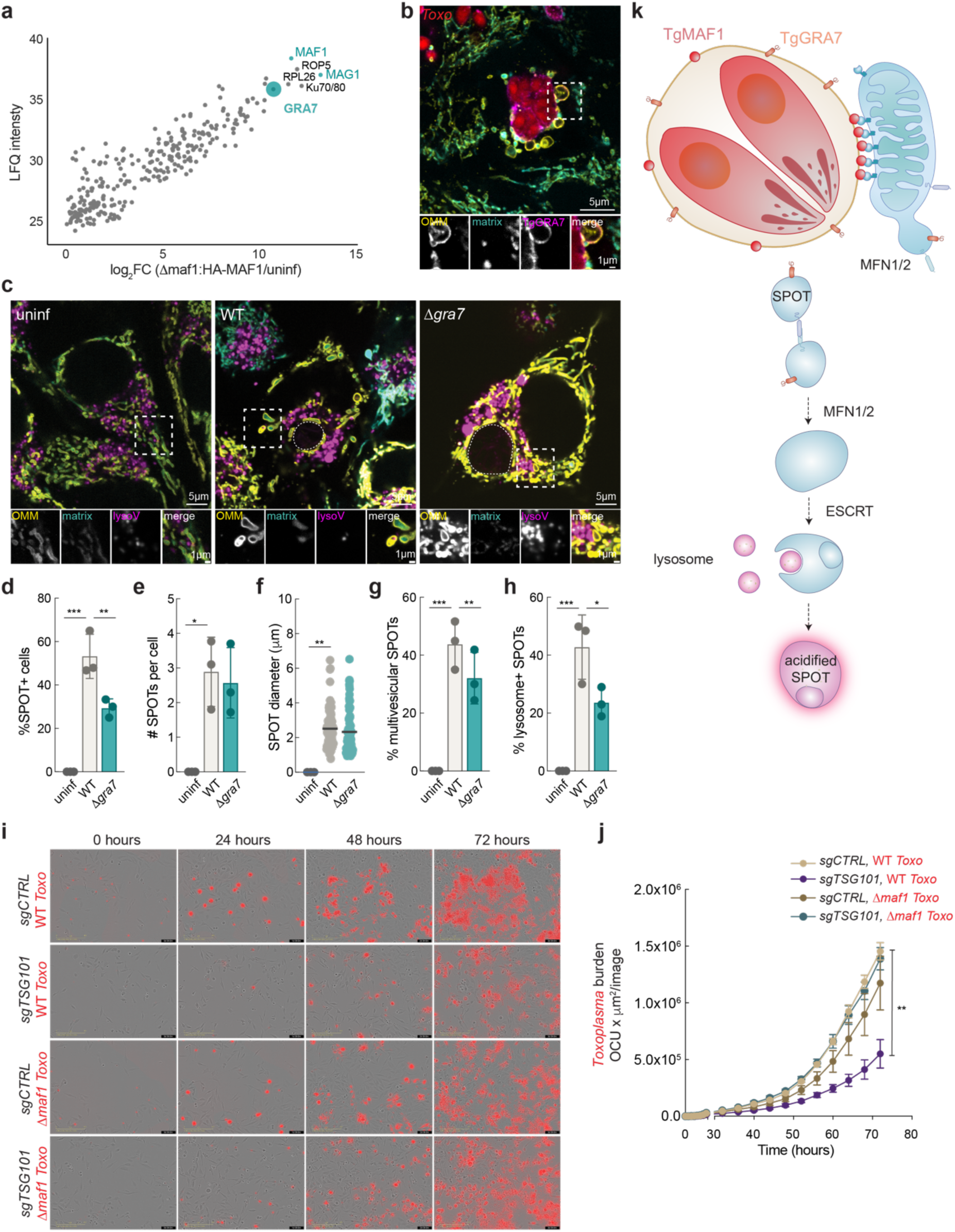
The secreted effector TgGRA7 promotes SPOT engulfment of lysosomes. (**a**) Anti-HA immunoprecipitates (IPs) from cells that were mock-infected (uninf) or infected with *Δmaf1*:HA-MAF1 parasites at an MOI of 1, 2.4, and 6 and analyzed by means of mass spectrometry; data for MOI 1/uninf are shown for that had a positive log_2_FC for the comparisons: all MOIs/uninf, MOI:6/MOI:2.4, MOI:6/MOI:1, MOI:2.4/MOI:1. LFQ, label-free quantification, TgMAF1 and most enriched dense granule interactors in teal (**b**) Representative immunofluorescence (IF) images of *Toxo*-infected SPOT reporter Hela cells at 24 hpi. (inset) TgGRA7 localizes to SPOTs associated at the PVM. Scale bars 5μm and (inset) 1μm. (**c**) Representative live-cell images of SPOT reporter infected with WT and *Δgra7 Toxoplasma-*parasites and labeled with lysoview at 24 hpi; scale bars 5 μm and inset 1 μm. (**d**) Percentage (%) of SPOT-positive cells, (**e**) number (#) SPOTs per cell, (**f**) SPOT diameter, (**g**) % of multivesicular SPOTs, and (**h**) % of lysoview-positive SPOTs from experiments as in (C); data are mean ± SEM from more than 30 infected cells from three replicates; *p<0.05; **p<0.01; ****p<0.0001 for uninf versus infected by means of one-way ANOVA analysis. (**i**) Representative IncuCyte images of sgCTRL and sgTSG101 HeLa cells infected with WT and *Δmaf1* parasites, rinsed at 2 hours after infection, and analyzed at 24 hour intervals after infection by means of IncuCyte for *Toxoplasma.* (**j**) Analysis of *Toxoplasma* burden from images as in (i) using orange mean intensity (OCU) / mm^2^ per image. Data are mean ± SEM from three replicates, **p< 0.01 by means of two-way ANOVA analysis. (**k**) Cartoon schematic of SPOT maturation.

TgGRA7 was readily observed at SPOTs emerging from the mitochondria in contact with the PVM unlike other effectors including TgMAF1 and TgMAG1, which were restricted to the PVM and PV, respectively (**Fig. 5b, Extended Data Fig. 14**). To address whether TgGRA7 influenced SPOT maturation, we analyzed SPOTs in cells infected with *Δgra7* parasites ^30^. The loss of TgGRA7 significantly impaired the formation rate and development of SPOTs in infected cells, despite SPOT number and size being unaffected (**Fig. 5c-h**). Consistent with this finding, the treatment of infected cells with pyrimethamine, which inhibited parasite replication and thus effector secretion, impaired SPOT maturation (**Extended Data Fig. 15**). Thus, TgGRA7 is targeted to nascent SPOTs and promotes their maturation.

### SPOTs promote *Toxoplasma* growth in an ESCRT-dependent manner

Our findings thus far established that active parasite secretion contributed to ESCRT-dependent maturation of SPOTs ^10^. We therefore asked whether SPOT maturation benefited *Toxoplasma.* To address this question, we compared *Toxoplasma* growth in cells depleted of TSG101. We found that the loss of TSG101 significantly reduced WT *Toxoplasma* burden by 50% at 72 hpi but had no effect on the growth of *Δmaf1* parasites that are deficient for SPOT formation (**Fig. 5i-j**). These results indicate that ESCRT machinery has no broad effect on overall *Toxoplasma* fitness, but rather supports parasite growth in the context of SPOT formation.

## Discussion

The ability of an intracellular pathogen to remodel existing host cell membranes serves several functions ranging from promoting invasion to evading destruction by host lysosomes ^1,2^. Our results demonstrate that, in addition to established mechanisms of remodeling existing membranes, the intracellular pathogen *Toxoplasma* drives mitochondria to shed SPOTs that acquire lysosomes and mature into acidified compartments in under twenty-four hours.

Although host ESCRT machinery was required for SPOT uptake of lysosomes, the canonical ESCRT recruitment factors PI3P and ubiquitylation were not. Instead, the parasite effector TgGRA7 mediated SPOT lysosome acquisition, indicating that *Toxoplasma* hijacks host ESCRT to drive SPOT maturation. The distribution of TgGRA7 to SPOTs raises the possibility that TgGRA7 recruits ESCRT machinery to SPOTs. Alternatively, TgGRA7 in conjunction with other effectors may induce membrane perturbations in SPOT-generating mitochondria that facilitate ESCRT recruitment, analogous to ESCRT recruitment to vesicles derived from the IMM that are subsequently targeted to lysosomes ^31^.

In line with SPOT maturation being a parasite-driven phenomenon, the depletion of host ESCRT machinery led to decreased *Toxoplasma* growth in wild-type parasites, but not in *Δmaf1* parasites that do not tether host mitochondria nor form SPOTs. These results suggest that the acidification of SPOTs may help counter a function of mitochondria or SPOTs that are induced in a TgMAF1-dependent manner. Given that MFN1/2-dependent elongation promotes the sequestration of fatty acids by these organelles, one possibility is that the acidification of SPOTs—which become enriched for MFN1 and MFN2 following their formation —may serve to prevent the sequestration of these nutrients within mitochondria or SPOTs ^32^.

SPOTs share several hallmarks of organelles: they are enclosed by multiple membranes, have a unique protein composition, acquire specific cargo through a regulated process, and develop a distinct biochemical environment, properties that together distinguish them from their mitochondrial progenitors and other organelles. Whether SPOTs are self-sustaining or heritable independent of infection remains an open question. Nevertheless, the degree of functional specialization exhibited by SPOTs suggests that the mitochondrial outer membrane can be reprogrammed toward organelle-like identity under pathological conditions.

In conclusion, our findings reveal a process by which a pathogen co-opts mitochondria-derived membranes to generate nascent organelles that support parasite growth yet are not themselves the replication niches of the parasite. More broadly, this work raises the possibility that infection can actively reshape the host organelle landscape in ways that expand the functional repertoire of the cell.

## Acknowledgments

We thank Ilian Atanassov and Xinping Li and the rest of the MPI-AGE Proteomics Core; Marcel Kirchner and the MPI-AGE FACS and Imaging Core for excellent microscopy support, Katrin Seidel and the CECAD imaging facility for electron microscopy support. We also thank Silvia Reato for excellent lab support, Tim Bartsch and Sebastian Kreimendahl for critical manuscript feedback, and all members of the Pernas lab for helpful discussions.

## Author contributions

Conceptualization: XL, JD, and LP; Methodology: all authors; Investigation: all authors; Funding acquisition: LP; Project administration: LP; Writing – original draft: XL, JD, LP; Writing – review & editing: All authors; Supervision: LP

## Data and materials availability

All data are available in the main text or the supplementary materials.

## List of Supplementary Materials

Table 1-2 (proteomics of *Toxo*-mito, cyto-mito and uninfected mito; and proteomics of TgMAF1Ps)

### Materials and Methods

#### Cell culture and cell lines

HeLa adenocarcinoma cells were obtained from ATCC (CCL-2); SPOT reporter HeLa cells stably expressing pMXs-3XHA-eGFP-OMP25 (referred to as OMM-GFP; Addgene #83356)(plasmid described in ^33^ and matrix-BFP (previously described ^10^; HeLa cells expressing pMXs-3xmyc-eGFP-OMP25 (Addgene # Addgene #83355)(plasmid described in ^33^; HeLa cells expressing pLenti mRFP-GFP (gift from C. Shoemaker, Dartmouth College) and OMM-BFP (previously described ^10^; HeLa cells expressing pLenti mRFP-eGFP-Fis1(gift from C. Shoemaker, Dartmouth College) were generated through lentiviral transduction. All cell lines were cultured in complete DMEM (cDMEM: DMEM and 10% heat-inactivated FBS). Cells were tested every 2 weeks for *Mycoplasma* infection by PCR.

#### Parasite culture and strains

*Toxoplasma gondii* parasites of the Type I (RH*Δhxgprt*) strain (deleted for the hypoxanthine-xanthine-guanine phosphoribosyl transferase (*HXGPRT*) gene), RH*Δku80:mCherry+*, and RH*Δku80ΔTgMAF1:mCherry+* (previously described ^11^), and RH*Δku80:mCherry+* parasites expressing HA, HA-MAF1 (previously described ^10^); Type I RHΔ*gra7* parasites were provided by Isabelle Coppens, (John Hopkins U) ^30^; Type I RHΔ*gra14* and Type I RHΔ*gra14:GRA14HA* parasites were provided by Dr. Vern Carruthers (U of Michigan, Ann Arbor) ^27^; were maintained by serial passage in human foreskin fibroblast (HFF) monolayers in cDMEM.

#### siRNA treatment

Cells were transfected ∼24-48 hours prior to infection (fresh siRNA readded during infection) using Lipofectamine RNAiMax (Invitrogen) according to manufacturer’s instructions. The following siRNAs were used:

*LUC (*CGUACGCGGAAUACUUCGA)TT

*MFN1* (GAUCGAAUAGCCACUGAAGAU)TT

*MFN2* (GCCUACCCUGUGAAGAUCUCU)TT

*OPA1* (GGUCUGCCAGUCUUUAGUGAA)TT

*STX17* (GUUGCAUGAAGAGCAUAUCAA)TT

RAB7 (GACCUCUAGGAAGAAAGUGUU)TT

*TSG101* (s14439) and *ALIX* (138251) were ordered from ThermoFisher and used according to the manufacturer’s instructions.

#### Plasmid descriptions

For transient expression, cells were transfected ∼12 hours prior to infection using X-tremeGene reagent (Sigma) per manufacturer’s instructions with CD63-peGFP (Addgene #62964), eGFP-LC3 (Addgene #11546), and mCherry-2xFYVE (Addgene #140060). For stable expression of OMM-targeted GFP, the triple hemagglutinin (3XHA-) and enhanced green fluorescent protein (eGFP) vector pMXs-3XHA-eGFP was used (Addgene #83356). For stable expression of OMM-targeted myc, the triple myc (3Xmyc-) and enhanced green fluorescent protein (eGFP) vector pMXs-3Xmyc-eGFP was used (Addgene #83355). For stable expression of OMM-targeted BFP, the 3XHA-eGFP in Addgene #83356 was replaced with BFP. For stable expression of OMM-targeted mRFP-eGFP, the Fis1 transmembrane domain sequence was introduced C-terminally of mRFP-eGFP (BsrGI site) of pLenti mRFP-eGFP (provided by C. Shoemaker, Dartmouth College).

#### Generation of CRISPR Knockdown (KD) cells

To generate *ALIX, TSG101, and HGS* CRISPR KD HeLa cells, the following sgRNAs were cloned into the pLenti CRISPRv2 (Addgene #5296), packaged into lentiviral particles, and used to transduce subconfluent cells SPOT reporter HeLa cells:

ALIX Sense: CACCG CGTCCGCTGGACAAGCACGA

ALIX Antisense: AAAC TCGTGCTTGTCCAGCGGACG C

TSG101 Sense: CACCG ATCCGCCATACCAGGCAACG

TSG101 Antisense: AAAC CGTTGCCTGGTATGGCGGAT C

HGS Sense: CACCG GTTCGGGGTGATGACCCGTA

HGS Antisense: AAAC TACGGGTCATCACCCCGAAC C

Following selection in 1-2 ug/ml puromycin, populations were passaged and seeded for indicated experiments at 5 days post-transduction.

#### Immunofluorescence Assays and Antibodies

For IF analysis, cells were plated and infected in a 24 well glass-bottom sensoplate, fixed at 24 hpi with *Toxoplasma* in 4% paraformaldehyde (fresh) in prewarmed cDMEM for 20 min at 37C, permeabilized for 20 min at RT with 0.2% Triton X-100, blocked in 3% Bovine serum albumin (BSA) for 30 min, incubated in 1:250 of primary Ab indicated in text O/N, rinsed 3X in 1X PBS, and maintained in 1X PBS until imaging. Images were taken using an Olympus IXplore SpinSR spinning disk confocal microscope. Primary Abs: LC3 (abcam #ab192890), calnexin (GeneTex #GTX109669-100), Golgin-97 (CST #13192S), Sec24B (CST #12042), catalase (CST #12980), EEA1 (CST #2411), LBPA (EMB Millipore MABT837), GAPDH (CST #2118), LAMP2 (SCBT sc-18822), ATP6V0D1 (abcam #ab202899) were used at 1:250 or 1:500 O/N. Secondary Abs: Alexa Fluor Plus 405, Alexa Fluor Plus 488, Alexa Fluor Plus 594, Alexa Fluo Plus 647 (Thermo Fisher) were used at 1:1000. All images were taken with a 100X objective and excitation with 405, 488, 561 and 640 laser lines, and processed via cellSens software and ImageJ (FIJI).

#### Immunoblotting and antibodies

Whole cells were harvested in chilled lysis buffer (50mM Hepes-KOH pH 7.4, 40mM NaCl, 2mM EDTA, 1.5mM NaVO4, 50mM NaF, 10mM NaPyrophosphate, 10mM, Sodium-β-Glycerophosphate (disodium salt pentahydrate), 1% Triton X-100) and lysed for 30 min on ice. Lysates were subsequently centrifuged at 10 min at 14,000 x g at 4°C and the supernatant was transferred into a fresh tube with 5X SDS added to a final of 1X SDS. Following SDS-PAGE and gel transfer, membranes were blocked with TBS-0.05% Tween 20 (TBS-T) and 5% BSA for overnight in primary antibodies. Following incubation, blots were washed three times in PBS-T and then incubated with horseradish peroxidase (HRP)-conjugated anti-mouse IgG (CST #7076) or anti-rabbit IgG (CST #7074) at a 1:4000 dilution for 45 minutes and developed using a chemiluminescence system (Pierce ECL substrate or Pierce ECL Plus Substrate; ThermoFisher Scientific). The following antibodies were used: MFN1/2 (abcam ab57602), OPA1 (BD Biosciences 612606), LC3 (abcam ab192890), CS (CST #14309), ACTB (CST #4970), TOMM40 (Sigma HPA036231), TOMM70 (HPA048020), GAPDH (CST #2118), TSG101 (Proteintech 28283), ALIX (Proteintech 12422), LAMP2 (SCBT sc-18822), RAB7 (CST #9367), STX17 (abcam ab229646), p62 (Proteintech 66184), TUBA (Proteintech 66031), TgMAF1 (23), TgGAP45 (Dr. D Soldati, U. of Geneva), TOMM40 (Santa Cruz sc365467), we thank Dr. Y Nishikawa (National Research Center for Protozoan Diseases, Obihiro University, Hokkaido, Japan) for providing us with TgGRA14 antibody ^34^.

#### Flow Cytometry

To assess *Toxoplasma* burden, monolayers of HeLa cells were rinsed infected with mCherry-expressing parasites at an MOI:1, rinsed at 2 hpi to remove non-invaders, and replenished with fresh media. After indicated times, cells were washed with 1X PBS, trypsinized and fixed in prewarmed 4% paraformaldehyde in 1X PBS for 10 min. After a brief spin, cells were resuspended in 1X PBS, filtered through a 35 µm nylon mesh, and analysed on a Sony SH800S for mCherry mean fluorescence intensity (mFI). ∼10,000 infected cells were recorded for analysis per replicate. Flow cytometry data was analysed using software BD Biosciences FlowJo v10.10.0. Statistical analysis was performed on mFI values using GraphPad Prism 10.

#### Incucyte analysis of parasite growth

To assess parasite growth rates, we used the Sartorius Incucyte SX5 system. The Incucyte SX5 is housed in a humidified 37°C incubator with 5% CO_2_. Host cells were seeded in a Greiner 96-well glass bottom black dish at 0.1 x 10^5^ cells per well the day prior to infection. Adhered cells were infected with mCherry-expressing parasites at an MOI: 0.5, rinsed at 2hpi to remove non-invaders, replenished with fresh media, and then housed in the Incucyte SX5 for the 72h duration of the experiment. A Breathe-Easy membrane film was applied to the plate to safeguard potential spills of pathogen-infected biological samples. Three images per well were acquired using the 20X objective for phase and orange channels every 4h for the 72h duration. Images that were out of focus, non-fluorescent (uninfected), or of a sub-confluent area of the well were excluded in the analysis. Automated image analysis using the Basic Analyzer mode was used and optimized thresholding to remove detection of background fluorescence.

#### Lysosomal labelling and drug treatments

To image proteolytically active lysosomes, Magic Red Fluorescent Cathepsin B Assay (abcam) was used according to manufacturer’s instructions. To visualize acidified lysosomes, infected cells were incubated with LysoTracker Red and Deep Red (ThermoFischer Scientific) at 50 nM or LysoView (Biotium) at a 1:1000 dilution in prewarmed cDMEM for 30 minutes prior to imaging. To inhibit global ubiquitylation, infected cells were treated with TAK-243 (1 μM) at 2 hours post infection (2 hpi). To inhibit PI3P synthesis, infected cells were treated with SAR-405 (10μm) at 2 hpi.

#### Quantification of lysosomes

Fluorescence microscopy images were analyzed using Fiji (ImageJ). LysoView 633 pH sensitive probe was used to label active lysosomes. Images were processed by applying a threshold and manually drawing regions of interests (ROI) that correspond to individual cells. Fluorescence signal from the parasite vacuole was excluded from the analysis using a binary mask. The Analyze Particles tools was used to quantify lysosomes in the ROI with 0.1-2 micron size and exclude signal on edges as parameters.

#### Mitotracker labeling

For staining with MitoTracker (Invitrogen), cells (24 hpi mock- or *Toxoplasma* infection) were incubated with prewarmed DMEM containing MitoTracker Deep Red at a concentration of 25 nM After 20 min of incubation at 37°C, cells were rinsed with prewarmed 1X PBS and then incubated in prewarmed cDMEM.

#### Isolation of Toxo-mito and cyto-mito

All of the following steps were performed on ice or at 4 °C. Equipment included a magnetic tube holder (Invitrogen DynaMag), a refrigerated centrifuge, and a vortex adaptor compatible with 1.5 mL microcentrifuge tubes. A 1 mL syringe fitted with a 27 G ¾” needle was used for mechanical lysis. HA- and Myc-tag antibody-coupled beads (Thermo Scientific #88837 and #88842) were pre-equilibrated by washing 150 μL per sample of HA bead mix three times in 1 mL of cold 1× PBS. Washed beads were resuspended in 100 μL 1× PBS per sample and aliquoted into pre-labeled 1.5 mL microcentrifuge tubes. 3XMyc-eGFP-OMP25 expressing HeLa cells were seeded in 10 cm dishes at a density of 2–10 × 10⁶ cells per dish. On the following day, cells were infected with *Δmaf1*:HA-MAF1-mCherry *Toxoplasma* parasites at a multiplicity of infection (MOI) of 6 and incubated for 8–12 hours prior to lysis. Media was aspirated, and cells were rinsed twice rapidly with ice-cold 1× PBS. Cells were scraped into 1 mL ice-cold 1× PBS containing protease and phosphatase inhibitors and transferred into a prechilled 2 mL tube. Mechanical disruption was achieved by passing the cell suspension through a 27 G syringe 10 times while avoiding the formation of bubble. A 25 μL aliquot was saved as a whole cell lysate (WC) input control (for immunoblot analysis). Remaining lysate was clarified by centrifugation at 700 × g for 2 min at 4 °C. The cleared supernatant was transferred to HA-tag antibody-coated beads and incubated with gentle rotation for 10 min at 4 °C. Tubes were positioned such that the cap was in contact with the rotator surface to minimize bead displacement. Following incubation, beads were pelleted on a magnetic rack for 1 min and the supernatant was collected and transferred to Myc-tag antibody-coated beads for a second round of immunoprecipitation for 5–10 min at 4 °C. Beads from each immunoprecipitation were washed twice with 1 mL of cold 1× PBS containing inhibitors, with 1 min incubations on the magnetic stand. The third wash step included transfer of beads to fresh, pre-labeled tubes. Beads were then resuspended in 80 μL of lysis buffer supplemented with inhibitors and incubated on ice for 20 min with intermittent vortexing. Lysates were cleared by centrifugation (14,000 × g, 10 min, 4 °C), and supernatants were transferred to clean tubes as HA- or Myc-IP samples.

#### Proteomics sample preparation and TMTpro labeling

Proteomics sample preparation was carried out as previously described ^35^. After washing the beads three to four times with detergent-free buffer (50 mM Tris-HCl, pH 7.5), 20–25 µL of lysis buffer (6 M guanidinium chloride (GuCl), 100 mM Tris-HCl, 2.5 mM TCEP, 10 mM chloroacetamide (CAA)) was added to the beads. The bead suspensions were heated at 95 °C for 10 min, the supernatants were collected and diluted 10-fold with 20 mM Tris-HCl, and the diluted lysates were sonicated with a Bioruptor (Diagenode) for 10 min (10 cycles). Subsequently, 100 ng of trypsin was added to each sample and proteins were digested at 37 °C overnight. The digestion was stopped the next morning by adding formic acid to a final concentration of 1%. Protein digests were then desalted using home-made C18 STAGE tips. Four micrograms of the peptides were dried and reconstituted in 9 µL of 0.1 M TEAB. Tandem mass tag (TMTpro, Thermo Fisher Scientific, cat. no. A44522) labeling was carried out according to the manufacturer’s instructions with the following changes: 0.5 mg of TMTpro reagent was resuspended in 33 µL of anhydrous ACN. Seven microliters of TMTpro reagent in ACN were added to 9 µL of clean peptide in 0.1 M TEAB. The final ACN concentration was 43.75% and the ratio of peptides to TMTpro reagent was 1:20. After 60 min of incubation, the reaction was quenched with 2 µL of 5% hydroxylamine. Labelled peptides were pooled, dried, resuspended in 200 µL of 0.1% formic acid (FA), split into two equal parts, and desalted using home-made C18 STAGE tips. One of the two parts was fractionated on a 1 mm × 150 mm ACQUITY column packed with 130 Å, 1.7 µm C18 particles (Waters, cat. no. SKU: 186006935), using an Ultimate 3000 UHPLC (Thermo Fisher Scientific). Peptides were separated at a flow of 30 µL/min with an 88 min segmented gradient from 1% to 50% buffer B for 85 min and from 50% to 95% buffer B for 3 min; buffer A was 5% ACN, 10 mM ammonium bicarbonate (ABC), and buffer B was 80% ACN, 10 mM ABC. Fractions were collected every three minutes, pooled in two passes (1 + 17, 2 + 18, etc.), and dried in a vacuum centrifuge (Eppendorf).

#### Proteomics LC-MS/MS analysis

Dried fractions were resuspended in 0.1% formic acid (FA) and separated on a 50 cm, 75 µm Acclaim PepMap column (Thermo Fisher Scientific, product no. 164942) and analysed on an Orbitrap Lumos Tribrid mass spectrometer (Thermo Fisher Scientific) equipped with a FAIMS device (Thermo Fisher Scientific). The FAIMS device was operated with two compensation voltages, -50 V and -70 V. Synchronous precursor selection (SPS)-based MS3 was used for the acquisition of the TMTpro reporter ion signals. Peptide separations were performed on an EASY-nLC 1200 using a 90 min linear gradient from 6% to 31% buffer B; buffer A was 0.1% FA and buffer B was 0.1% FA, 80% ACN. The analytical column was operated at 50 °C. Raw files were split based on the FAIMS compensation voltage using FreeStyle (Thermo Fisher Scientific).

#### Proteomics data analysis

Proteomics data was analyzed using MaxQuant, version 1.6.17.0, ^36^. Peptide fragmentation spectra were searched against the reviewed and unreviewed sequences of the human reference proteome (proteome ID UP000000589, downloaded December 2018 from UniProt). Methionine oxidation and protein N-terminal acetylation were set as variable modifications; cysteine carbamidomethylation was set as fixed modification. The digestion parameters were set to “specific” and “Trypsin/P,” The minimum number of peptides and razor peptides for protein identification was 1; the minimum number of unique peptides was 0. Protein identification was performed at a peptide spectrum matches and protein false discovery rate of 0.01. The isotope purity correction factors, provided by the manufacturer, were included in the analysis. Differential expression analysis was performed using limma, version 3.34.9, ^37^ in R, version 3.4.3 ^38^. GO and KEGG pathway enrichment analyses on differentially expressed genes were performed in R v4.4 using Bioconductor packages, clusterProfiler v3.20. GO Biological Process enrichment was performed using enrichGO and KEGG enrichment analysis was performed using enrichKEGG.

#### Data availability

Data are available via ProteomeXchange with identifier PXD072097; token: Urcb5yrDz0fD

#### CLEM Imaging

Cells were grown in glass bottom dishes (MatTek, # P356-1.5-14-C) which were coated with a carbon finder pattern using a mask (Leica, # 16770162) and a carbon coater ACE 200 (Leica). Cells were fixed for 15 min at room temperature in 2% glutaraldehyde (Sigma, # G5882-100ML) with 2.5 % sucrose (Roth, # 4621.1) and 3mM CaCl_2_ (Sigma, # C7902-500G) in 0.1M HEPES buffer pH 7.4 (Roth, # 9105.1). Cells were washed three times with 0.1M HEPES buffer and fluorescent and brightfield images were taken using the Olympus IXplore SpinSR spinning disk confocal with the 60x oil objective. Localization coordinates of cells of interest were noted. Cells were incubated with 1% Osmiumtetroxid (Science Services, # E19190) and 1.5% Potassium ferricyanid (Sigma, # P8131) for 30 min at 4°C. After 3x5min wash with 0.1M Cacodylate buffer (Applichem, # A2140,0100), samples were dehydrated using ascending ethanol series (50%, 70%, 90%, 100%) (VWR, # 153386F) for 7 min each at 4°C. Cells were infiltrated with a mixture of 50% Epon/ethanol for 1 h, 66% Epon/ethanol for 2 h and with pure Epon (Sigma, # 45359-1EA-F) overnight at 4°C. TAAB capsules (Agar Scientific, Stansted, UK) filled with Epon were placed upside down onto the glass bottom and cured for 48 h at 60°C. Glass bottom was removed by alternatingly putting the dish into boiling water and liquid nitrogen. Block face was trimmed to the previous noted square using a 90° trimming tool (Diatome, Biel, Switzerland). Ultrathin sections of 70 nm were cut using an ultramicrotome (Leica Microsystems, UC6) and a diamond knife (Science Services # DU3530), collected onto pioloform (Plano, #R1275) coated slot grids (Science Service # G2010-Cu) and stained with 1.5 % uranyl acetate (Agar Scientific, # R1260A) for 15 min at 37°C and 3% Reynolds lead citrate solution made from Lead (II) nitrate (Roth, # HN32.1) and tri-Sodium citrate dehydrate (Roth #4088.3) for 3 min. Images were acquired using a JEM-2100 Plus Transmission Electron Microscope (JEOL) operating at 80kV equipped with a OneView 4K camera (Gatan). Overlay of TEM and LM images were generated using the EC-CLEM plugin (Paul-Gilloteaux et al., 2017) for the software ICY.

**Extended Data Fig. 1:**
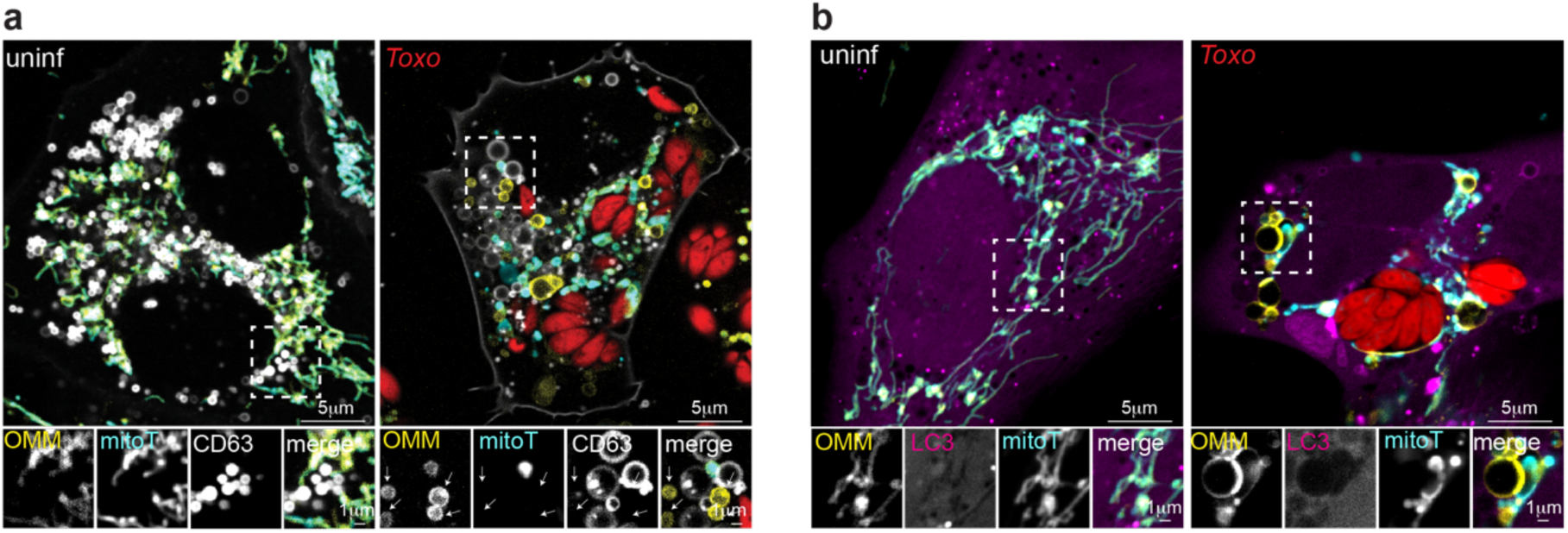
SPOTs are not trafficked to autophagosomes or multivesicular bodies. Representative live-cell imaging of (**a**) uninfected (uninf) and *Toxoplasma-*mCherry (*Toxo^mCh^*)-HeLa cells expressing OMM-BFP (yellow), CD63-GFP (white), were labeled with mitotracker (mitoT, cyan) and imaged at 24 hpi (**b**) OMM-BFP (yellow) and eGFP-LC3 (magenta) expressing HeLa cells were labeled with mitoT (cyan) at 24 hpi with *Toxo^mCh^*. Scale bars are 5 µm and 1 µm in the inset.

**Extended Data Fig. 2:**
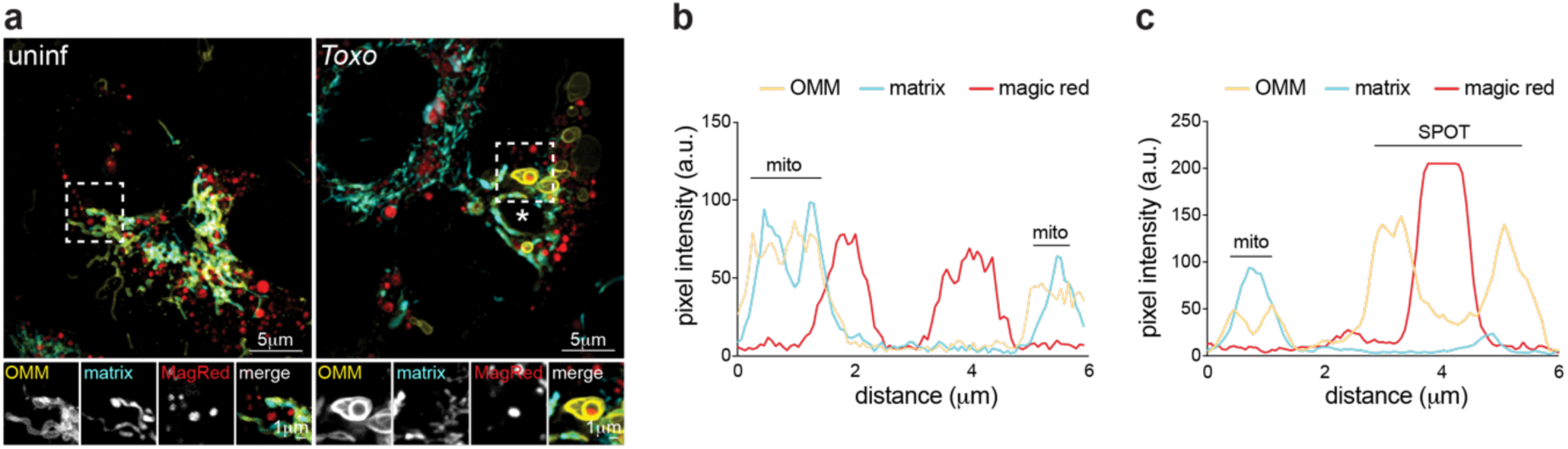
Lysosomes within SPOTs retain proteolytic activity. Representative live-cell imaging of SPOT reporter HeLa cells infected with *Toxoplasma* (white asterisk) incubated with a cell permeant Cathepsin B substrate (Magic Red Assay Kit) that fluoresces upon cleavage by Cathepsin B. Corresponding pixel intensity plots for white line in inset panels in (**a**): (**b**) uninfected and (**c**) infected.

**Extended Data Fig. 3:**
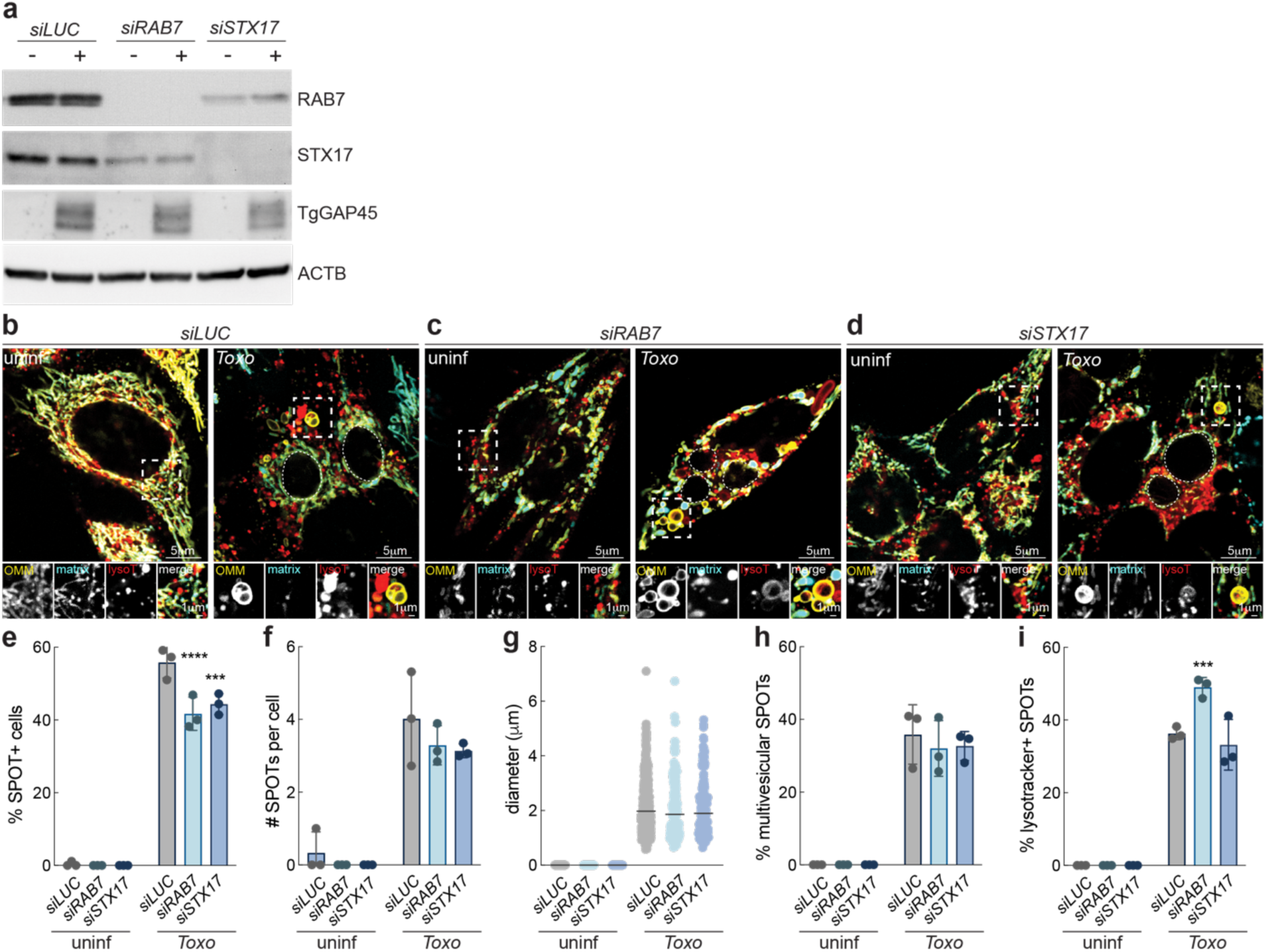
SPOT uptake lysosomes in an Rab7- and STX17- independent manner. **(a)** *CTRL, TSG101, ALIX, and TSG101 & ALIX-*silenced (si) SPOT reporter HeLa cells were analyzed by means of immunoblotting for RAB7 ∼23kDa; STX17 ∼35kDa; TgGAP45 ∼45kDa; and ACTB ∼45kDa. (**b-d**) Representative live-cell images of *CTRL, RAB7,* and *STX17-*silenced HeLa SPOT reporter cells infected with *Toxoplasma* (indicated by dotted lines) and labeled with lysotracker at 24 hpi; scale bars 5μm and inset 1μm. (**e**) Percentage (%) of SPOT-positive cells, (**f**) number (#) SPOTs per cell, (**g**) SPOT diameter, (**h**) % of multivesicular SPOTs, and (**i**) % of lysotacker-positive SPOTs from experiments as in (b-d); data are mean ± SEM from more than 30 infected cells from three replicates; ***p<0.001; ****p<0.0001 for *CTRL* versus silenced by means of two-way ANOVA analysis.

**Extended Data Fig. 4:**
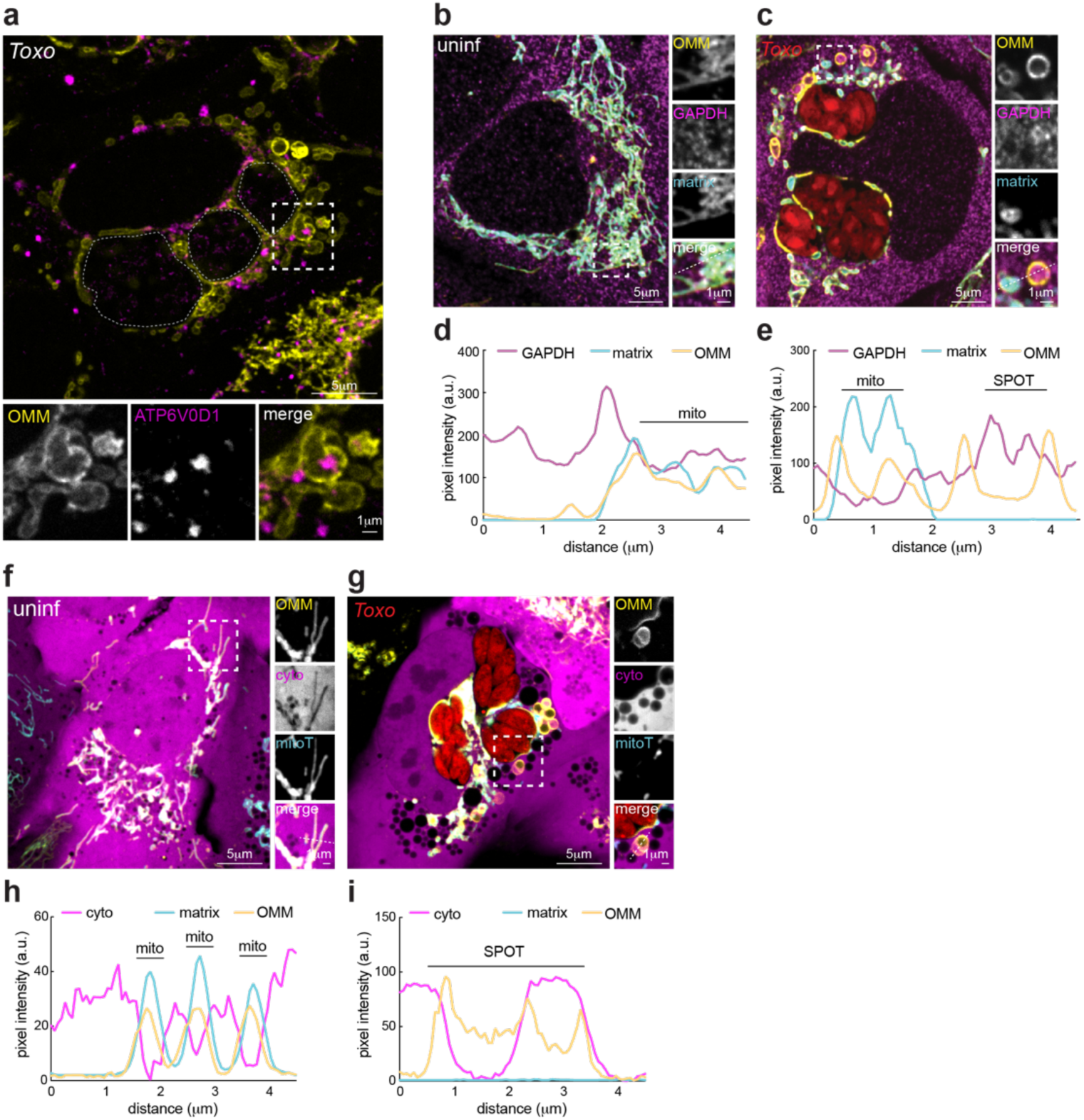
SPOTs engulf lysosomes and acquire cytosolic proteins. (**a**) Representative images of *Toxo-*infected SPOT reporter HeLa cells that were fixed at 24 hpi and processed for IF analysis of ATP6V0D1. Representative images of (**b**) uninfected and (**c**) *Toxo^mCh^*-infected HFFs expressing OMM-GFP and matrix-BFP that were fixed at 24 hpi and processed for IF analysis of cytosolic GAPDH. (**d** and **e**) Corresponding pixel intensity plots for white line in inset panels in (b and c). Representative live-cell images of (**f**) uninfected and (**g**) *Toxo^mCh^*-infected HeLa cells expressing OMM-BFP, GFP, and labeled with mitoTracker (mitoT) that were imaged at 24 hpi. (**h** -**i**) Corresponding pixel intensity plots for white line in inset panels in (f and g). Scale bars 5 μm and inset 1 μm.

**Extended Data Fig. 5:**
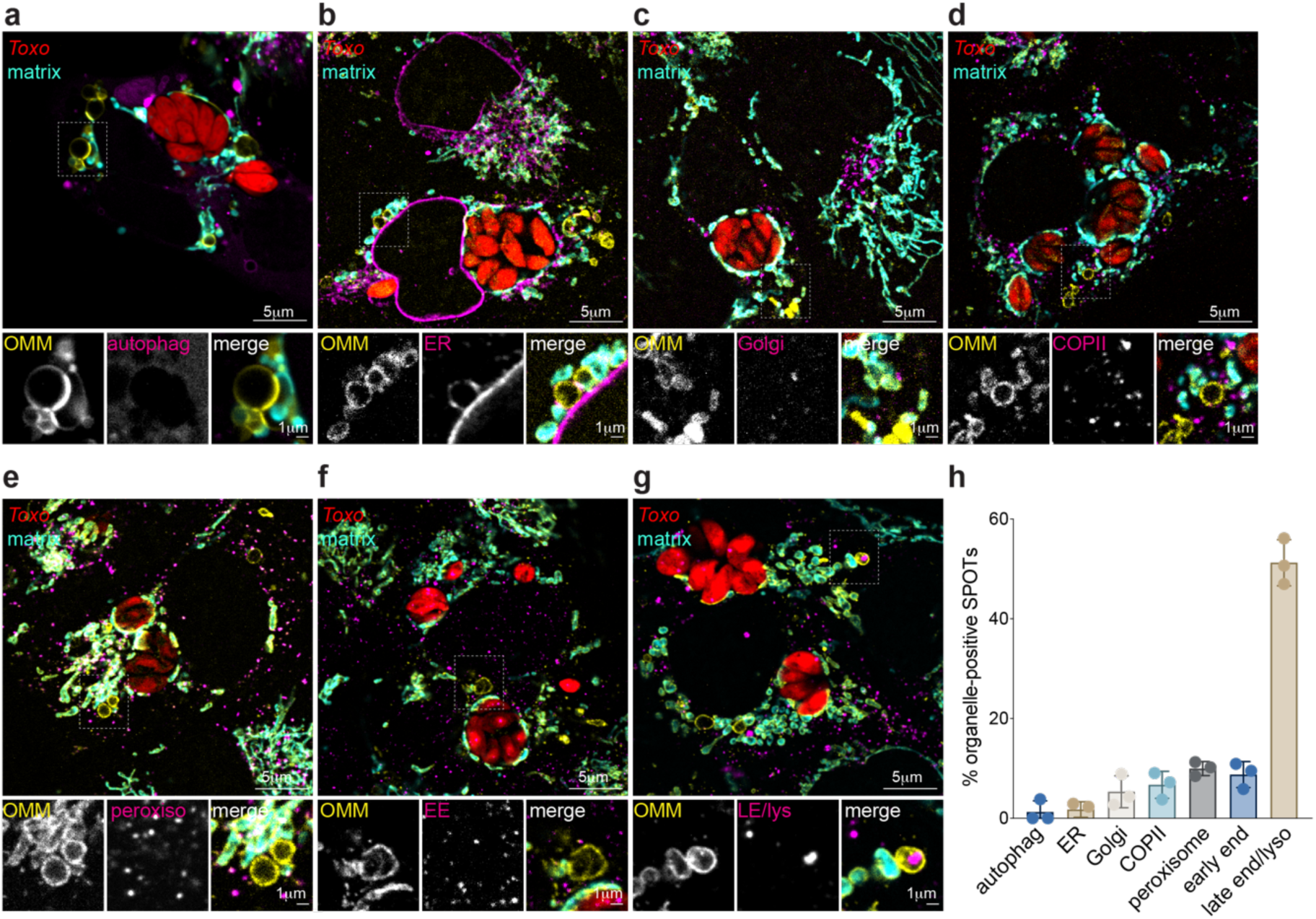
SPOTs selectively acquire lysosomes. Representative images of *Toxo^mCh^-*infected HFFs that were fixed at 24 hpi and processed for IF analysis of (**a**) autophagosomes, LC3; (**b**) endoplasmic reticulum (ER), calnexin; (**c**) Golgi, Golgin97 (**d**) COPII vesicles, Sec24B; (**e**) peroxisomes, catalase, (**f**) early endosomes, EEA1, and (**g**) late endosomes/lysosomes (LE/lys), LBPA; scale bar, 5 µm; inset, 1 µm. (**h**) % organelle-positive SPOTs from images as in (a-g); data are mean ± SEM from more than 30 infected cells from three replicates.

**Extended Data Fig. 6:**
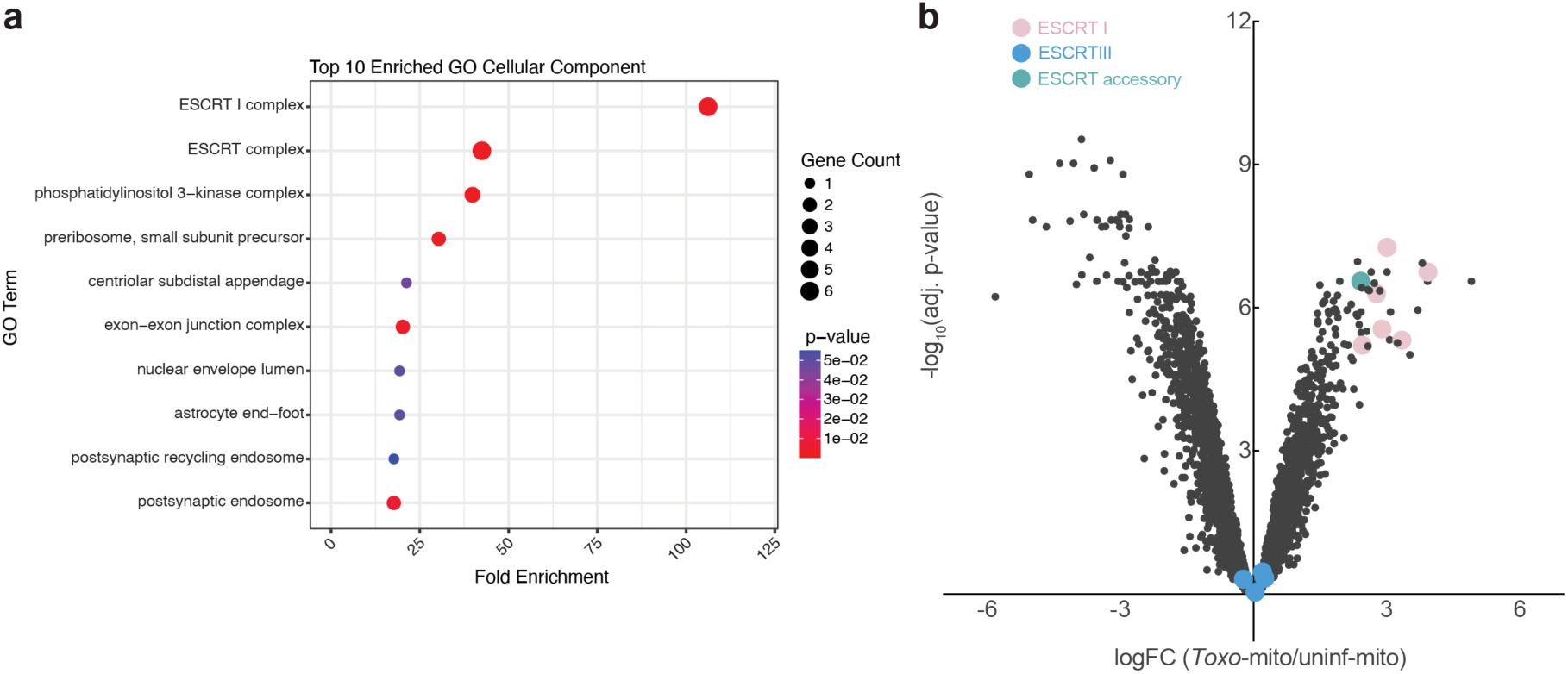
ESCRT-I components are enriched in *Toxo-*mito isolates. (**a**) GO pathway enrichment analyses on the top 100 differentially abundant proteins between *Toxo-*mito and uninf-mito from experiment performed as in (Fig. 3a-b) and analyzed by mass spectrometry; *Toxo*-mito (n=5) and uninf-mito (n=5) (**b**) Volcano plot of proteins identified in *Toxo*-mito and uninf-mito fractions. ESCRT-I proteins; pink; ESCRT-III proteins; blue; and ESCRT accessory proteins; teal enriched in *Toxo*-mito fraction are highlighted.

**Extended Data Fig. 7:**
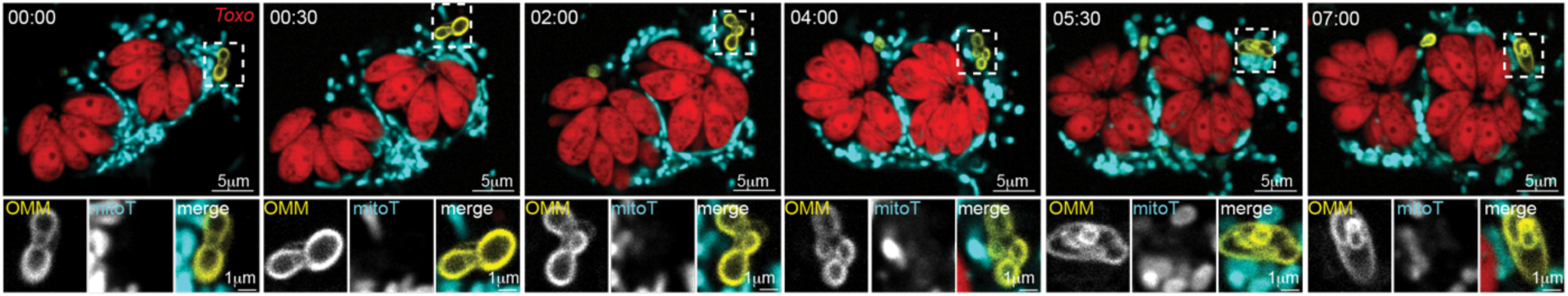
SPOTs accumulate intraluminal vesicles. A multivesicular SPOT was photoactivated in HeLa cells expressing OMM-targeted photo-activatable GFP that were infected with mcherry-expressing *Toxoplasma* and labeled with mitotracker and imaged over the indicated time frames (mm:hh). Scale bars 5μm and inset 1μm.

**Extended Data Fig. 8:**
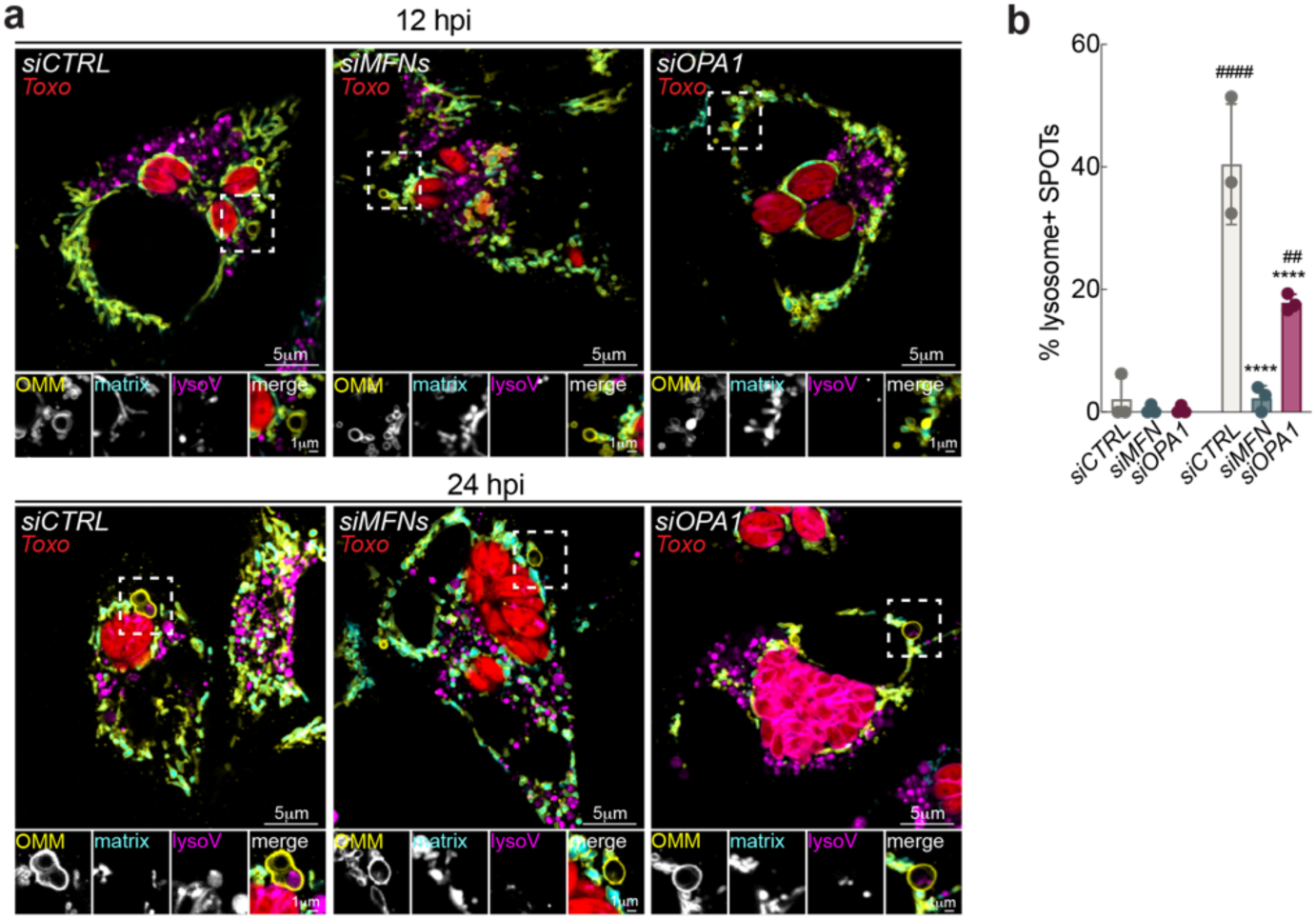
Loss of MFN1/2 impairs SPOT uptake of lysosomes. (**a**) Representative live-cell images of *CTRL, MFN1&2,* and *OPA1-*silenced (si) HeLa cells expressing OMM-targeted GFP and matrix-targeted BFP (SPOT reporter) labeled with lysotracker, infected with *Toxoplasma-*mCherry (*Toxo^mCh^*), and imaged at 12 hours post infection (hpi) and 24 hpi; scale bar 5μm, inset 1μm. (**b**) Percentage (%) of lysosome-positive SPOTs from images as in (a); data are mean ± SEM from more than 30 infected cells from three replicates; ## p<0.01, #### p<0.0001 for 12 hpi versus 24 hpi, ****p<0.0001 for *siCTRL* versus *siMFN, siOPA1* by means of two-way ANOVA analysis.

**Extended Data Fig. 9:**
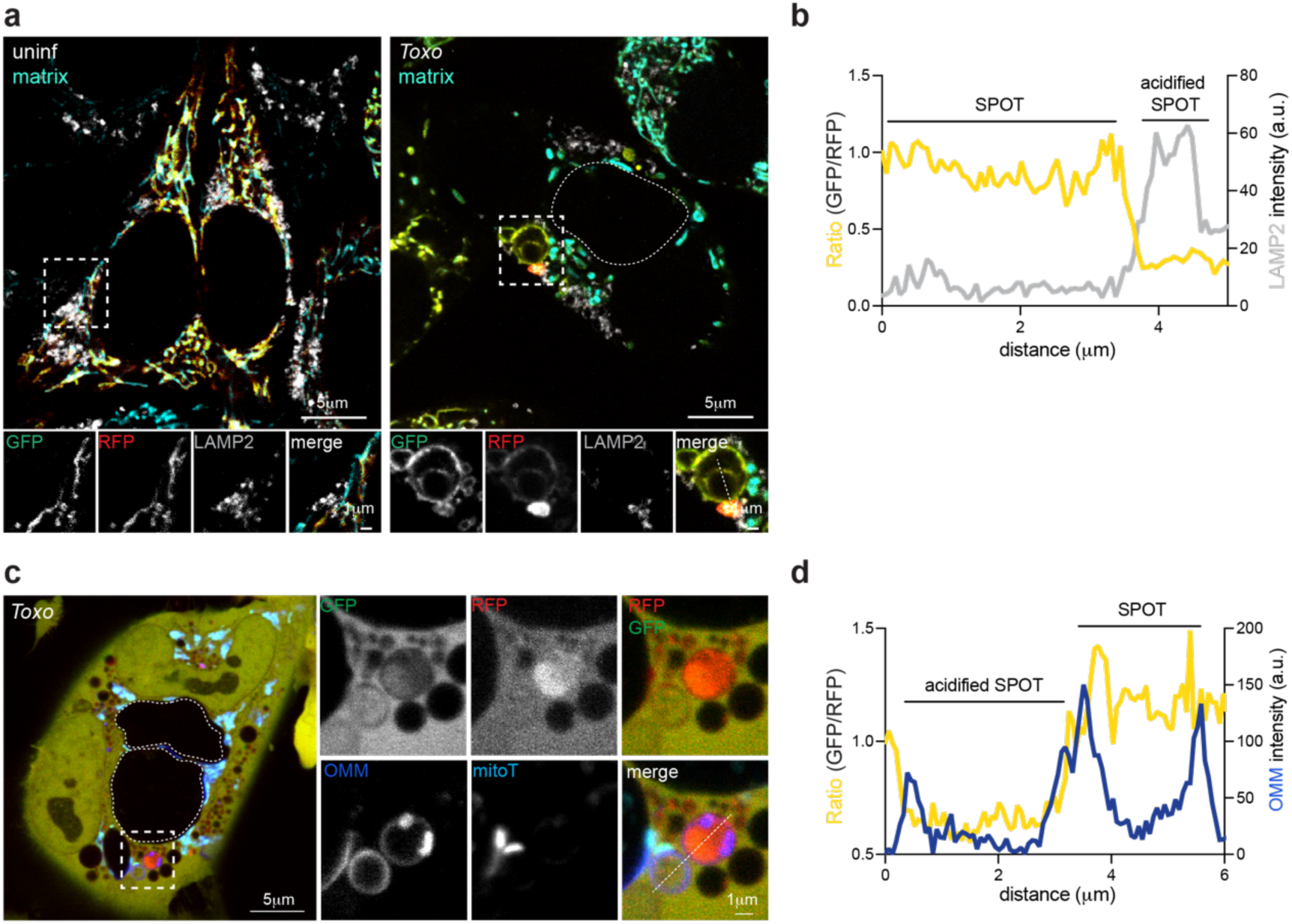
Lysosome-positive SPOTs become acidified. Representative images of (**a**) uninfected (left) and *Toxo*-infected (right) HeLa cells expressing an OMM-tandem fluorescent reporter (RFP-eGFP-Fis1^TMD^) of acidification that were fixed at 24 hpi and processed for IF analysis of LAMP2. (**b**) Corresponding pixel intensity plots for white line in inset panels in (a, *Toxo-*infected) going from top to bottom. Scale bars 5μm and inset 1μm. (**c**) Representative live-cell images of a *Toxo*-infected HeLa cells expressing OMM-BFP and a cytosolic tandem fluorescent reporter (RFP-eGFP) of acidification that was labeled with mitotracker at 24 hpi. (**d**) Corresponding pixel intensity plots for white line in inset panels in (c) going from top to bottom. Scale bars 5μm and inset 1μm.

**Extended Data Fig. 10:**
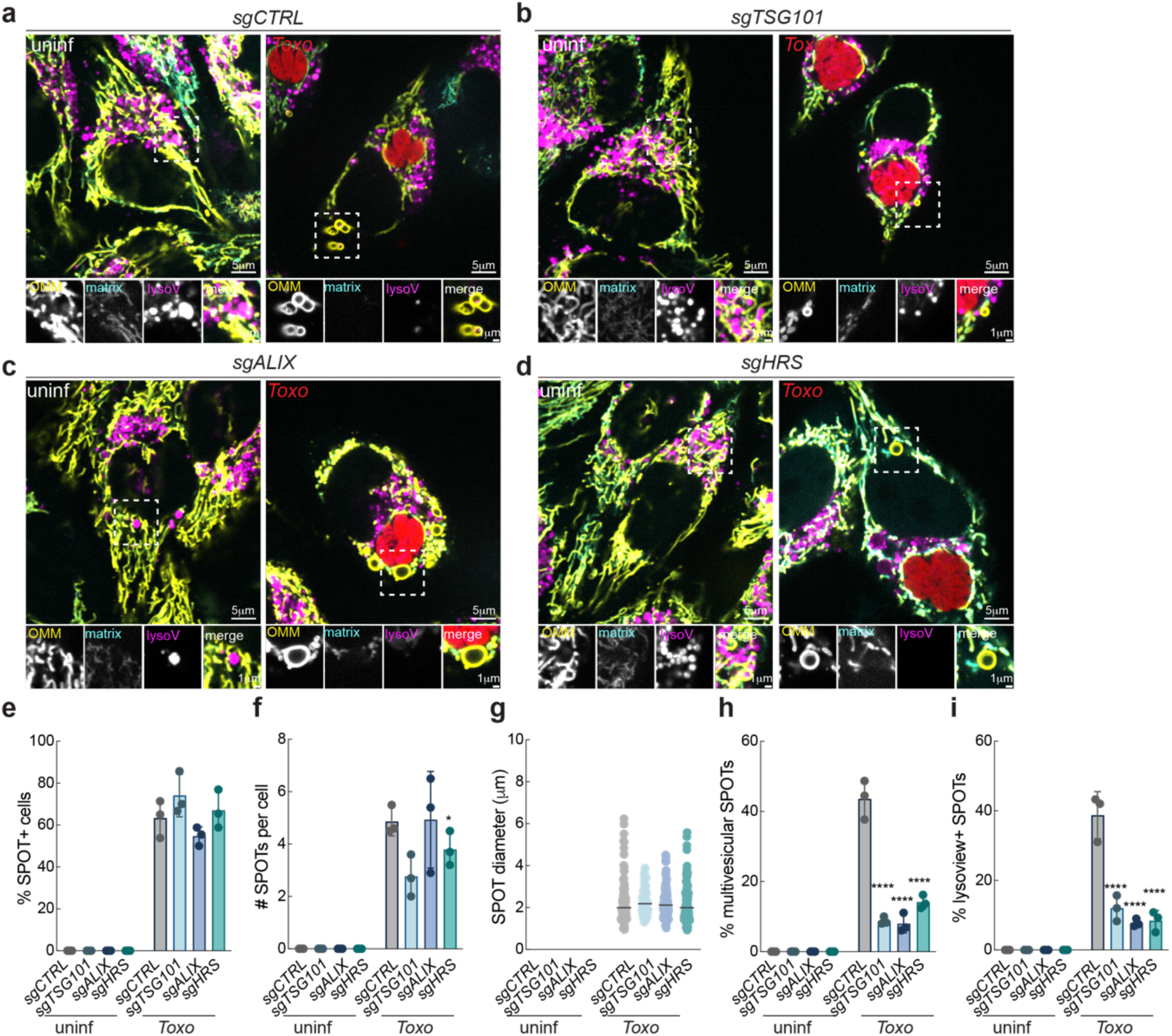
ESCRT-0 is required for SPOT maturation. Representative live-cell images of (**a**) *CTRL,* (**b**) *TSG101,* (**c**) *ALIX,* and (**d**) *HRS* CRISPR/Cas9-mediated knockdowns in SPOT reporter HeLa cells infected with *Toxoplasma-*mCherry (*Toxo^mCh^*) and labeled with lysoview at 24 hpi; scale bar 5μm, inset 1μm. (**e**) Percentage (%) of SPOT-positive cells, (**f**) number (#) SPOTs per cell, (**g**) SPOT diameter, (**h**) % of multivesicular SPOTs, and (**i**) % of lysoview-positive SPOTs from experiments as in (A-D); data are mean ± SEM from more than 30 infected cells from three replicates; *p<0.05, ****p<0.0001 for *CTRL* versus silenced by means of two-way ANOVA analysis.

**Extended Data Fig. 11:**
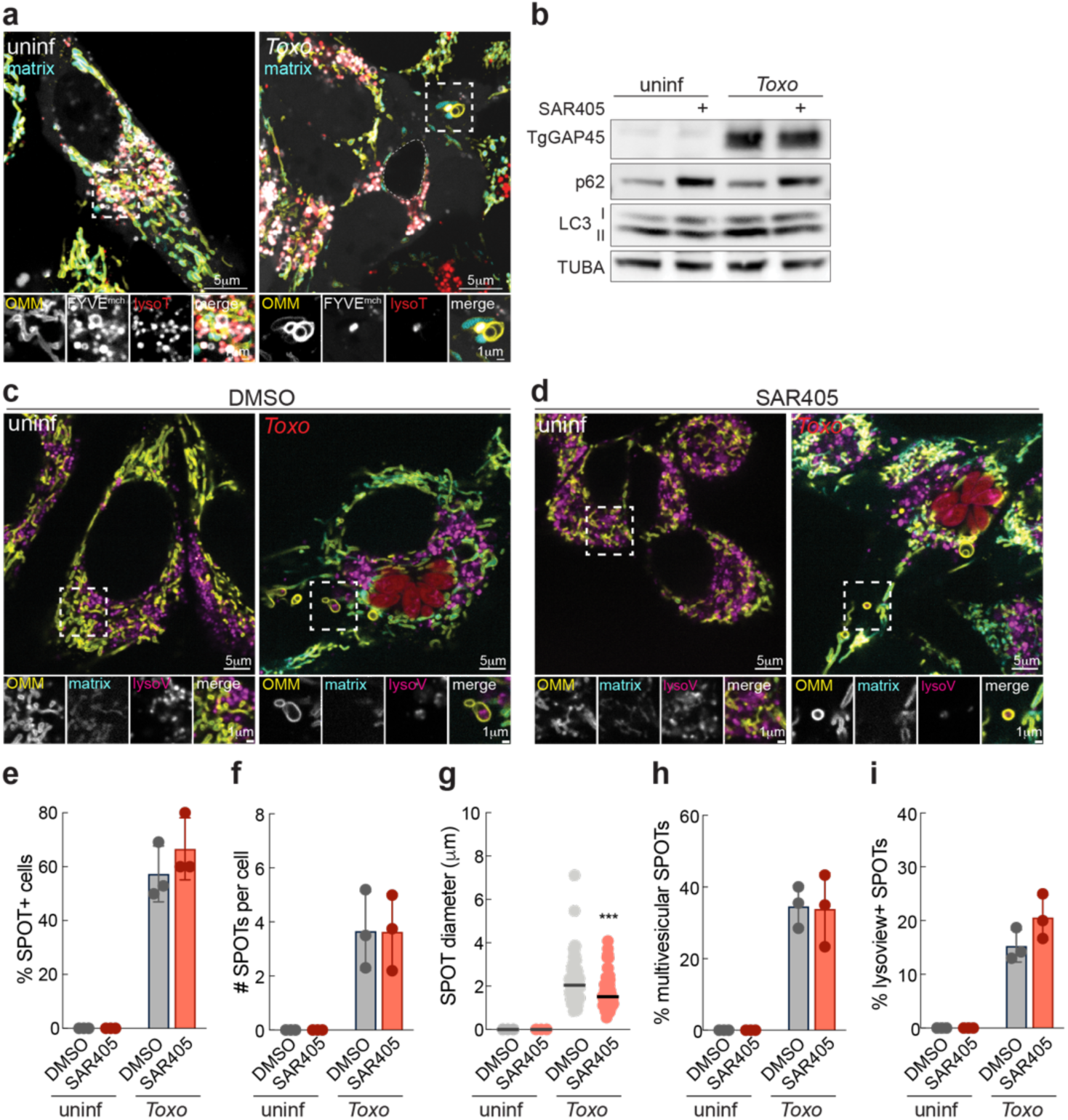
PI3P synthesis is dispensable for ESCRT-mediated maturation of SPOTs. (**a**) Representative live-cell images of uninfected and *Toxo*-infected SPOT reporter HeLa cells expressing 2XFYVE:mCherry and labelled with lysotracker at 24 hpi. Scale bars 5μm and inset 1μm. (**b**) HeLa cells were uninfected or infected with *Toxoplasma* and treated with SAR-405 at 2 hpi, harvested at 24 hpi, and analyzed by means of immunoblotting for TgGAP45 ∼45 kDa, p62 ∼62 kDa, LC3 ∼16-18 kDa, and TUBA ∼55kDa. Representative live-cell images of SPOT reporter HeLa cells that were uninf or infected with *Toxo^mCh^*, treated with (**c**) DMSO or (**d**) SAR-405 at 2 hpi, and labeled with lysotracker at 24 hpi; scale bar 5μm, inset 1μm. (**e)** Percentage (%) of SPOT-positive cells, (**f**) number (#) SPOTs per cell, (**g**) SPOT diameter, (**h**) % of multivesicular SPOTs, and (**i**) % of lysoview-positive SPOTs from experiments as in (c-d); data are mean ± SEM from more than 30 infected cells from three replicates; ***p<0.001 for DMSO versus SAR-405 by means of two-way ANOVA analysis.

**Extended Data Fig. 12:**
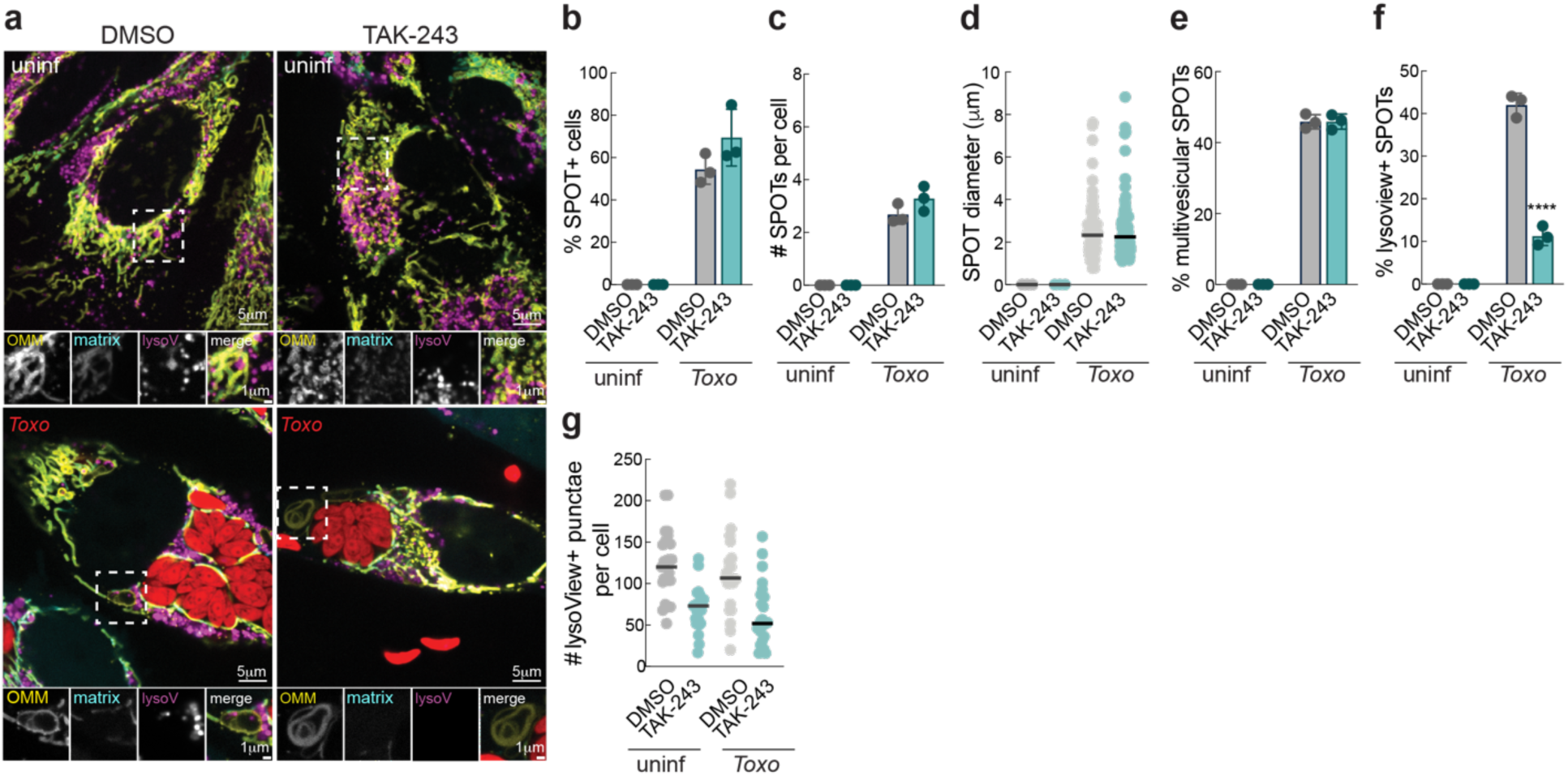
Ubiquitination is dispensable for ESCRT-mediated maturation of SPOTs. (**a**) Representative live-cell images of SPOT reporter HeLa cells that were uninf or infected with *Toxo^mCh^*, treated with DMSO or TAK-243 at 2 hpi, and labeled with lysotracker at 24 hpi; scale bar 5μm, inset 1μm. (**b**) Percentage (%) of SPOT-positive cells, (**c**) number (#) SPOTs per cell, (**d**) SPOT diameter, (**e**) % of multivesicular SPOTs, and (**f**) % of lysoview-positive SPOTs from experiments as in (A); data are mean ± SEM from more than 30 infected cells from three replicates; ****p<0.001 for *DMSO* versus TAK-243 by means of two-way ANOVA analysis. (**g**) Number (#) of lysoview-positive structures per cell, data are mean ± SEM from more than 25 cells from one replicate.

**Extended Data Fig. 13:**
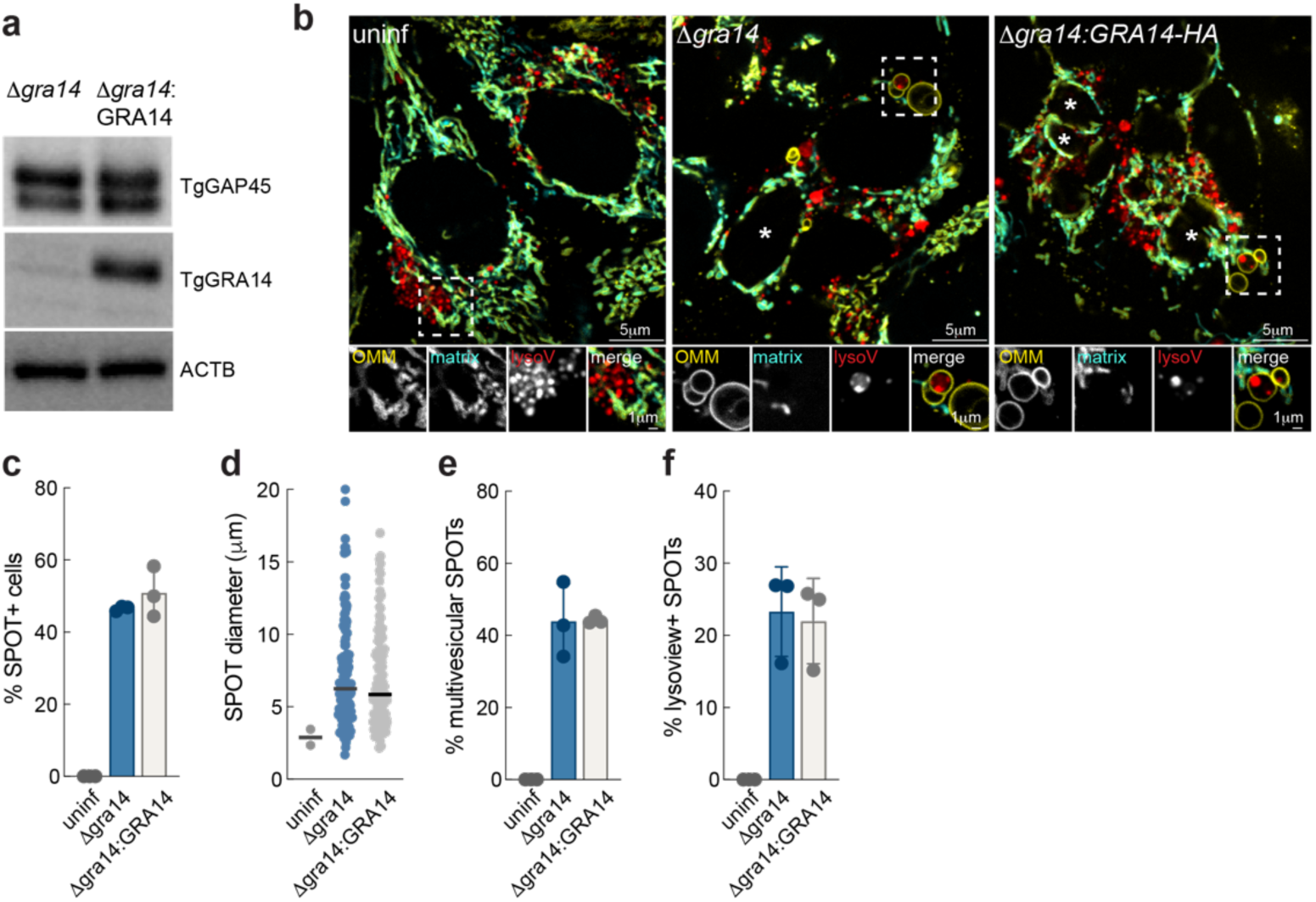
GRA14 is dispensable for SPOT uptake of lysosomes. (**a**) *Δgra14* and *Δgra14:GRA14_HA-*infected HeLa cells were analyzed by means of immunoblotting for TgGAP45 ∼45kDa; TgGRA14 (∼45kDa), and ACTB ∼45kDa. (**b**) Representative live-cell images of HeLa SPOT reporter cells uninfected or infected with *Δgra14* or *Δgra14::GRA14HA* parasites (indicated by asterisks), and labeled with lysoview (lysoV) at 24 hpi; scale bars 5μm and inset 1μm. (**c**) Percentage (%) of SPOT-positive cells, (**d**) SPOT diameter (μm), (**e**) % of multivesicular SPOTs, and (**f**) % of lysoV-positive SPOTs from experiments as in (B); data are mean ± SEM from more than 30 infected cells from three replicates.

**Extended Data Fig. 14:**
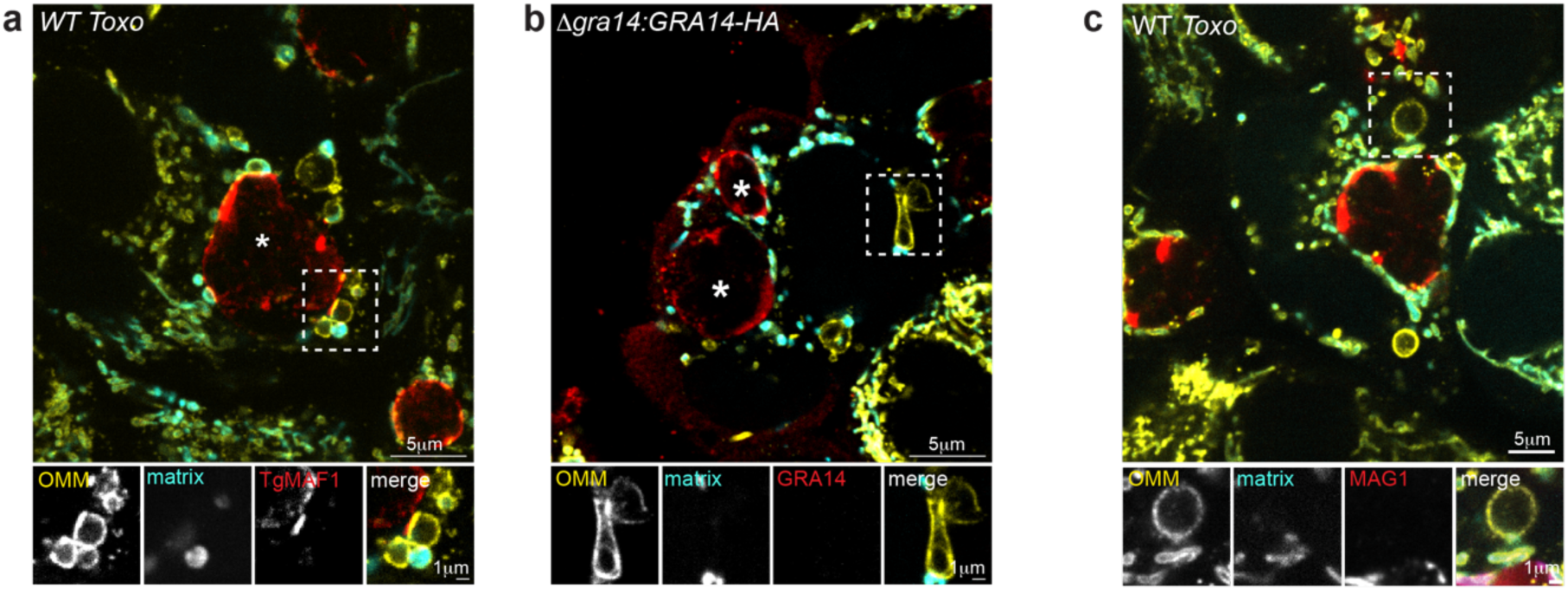
SPOTs lack TgMAF1, TgGRA14 and TgMAG1. Representative immunofluorescence (IF) images of SPOT reporter Hela cells infected with (**a**) WT *Toxoplasma* and probed for TgMAF1, (**b**) *Δgra14:GRA14-HA Toxoplasma* and probed for either TgGRA14, and (**c**) WT *Toxoplasma* and probed for TgMAG1. Scale bars 5μm and inset 1μm.

**Extended Data Fig. 15:**
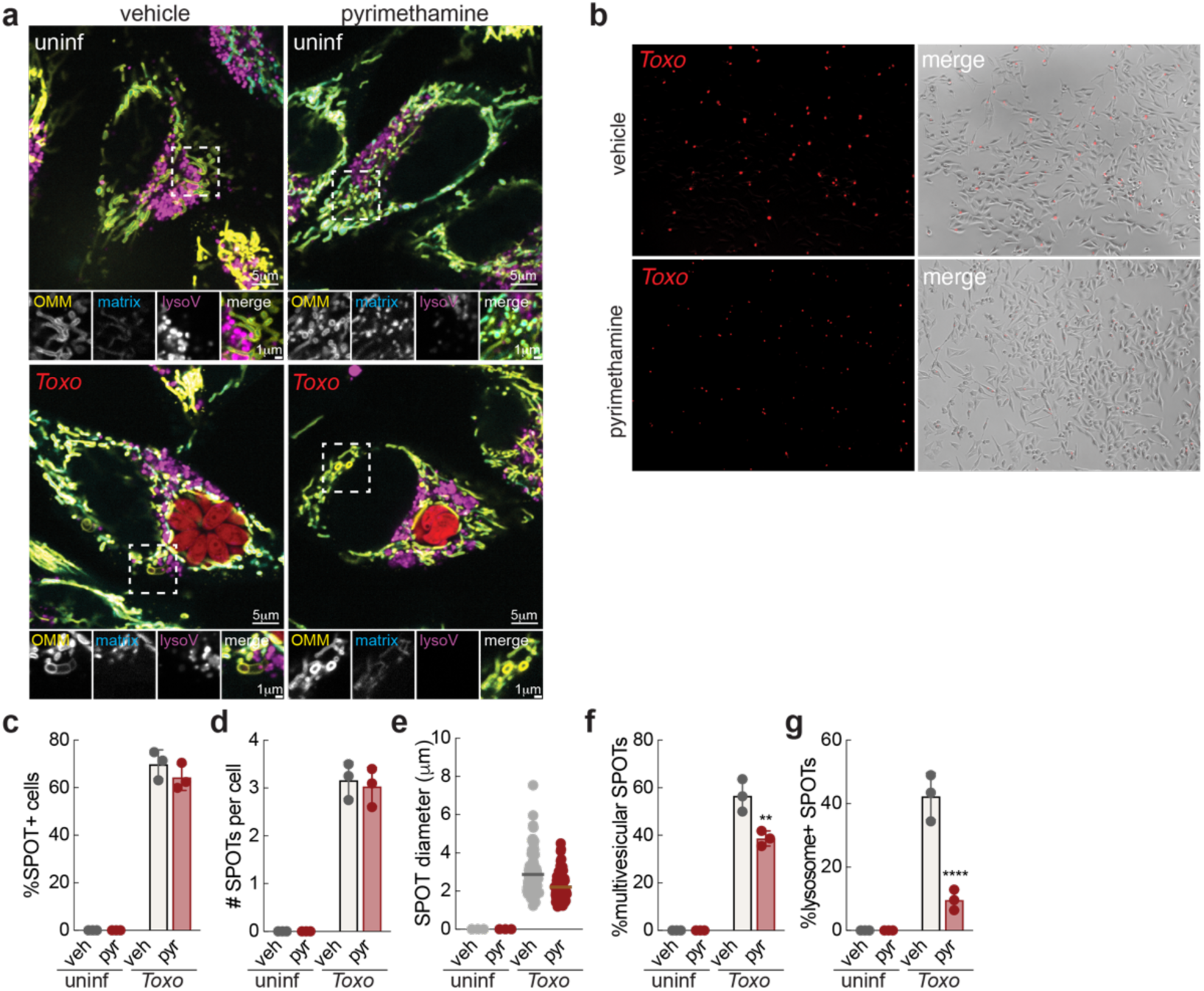
Inhibiting *Toxoplasma* growth impairs SPOT maturation. (**a** and **b**) Representative live-cell images of SPOT reporter HeLa cells that were uninf or infected with *Toxo^mCh^*, treated with vehicle (ethanol) or pyrimethamine (3μm) at 8 hpi, and labeled with lysotracker at 24 hpi; scale bar 5μm, inset 1μm. (**c**) Percentage (%) of SPOT-positive cells, (**d**) number (#) SPOTs per cell, (**e**) SPOT diameter, (**f**) % of multivesicular SPOTs, and (**f**) % of lysoview-positive SPOTs from experiments as in (a); data are mean ± SEM from more than 30 infected cells from three replicates; **p<0.001; and ****p<0.001 for vehicle versus pyrimethamine by means of two-way ANOVA analysis.

## References

1 Woida, P. J. & Lamason, R. L. Pathogen-induced rerouting of host membrane trafficking. Curr Opin Cell Biol 94, 102520 (2025). 10.1016/j.ceb.2025.102520

2 Cossart, P. & Roy, C. R. Manipulation of host membrane machinery by bacterial pathogens. Curr Opin Cell Biol 22, 547–554 (2010). 10.1016/j.ceb.2010.05.006

3 Pernas, L. & Scorrano, L. Mito-Morphosis: Mitochondrial Fusion, Fission, and Cristae Remodeling as Key Mediators of Cellular Function. Annu Rev Physiol 78, 505–531 (2016). 10.1146/annurev-physiol-021115-105011

4 Palade, G. E. An electron microscope study of the mitochondrial structure. J Histochem Cytochem 1, 188–211 (1953).

5 Neuspiel, M. et al. Cargo-selected transport from the mitochondria to peroxisomes is mediated by vesicular carriers. Curr Biol 18, 102–108 (2008). 10.1016/j.cub.2007.12.038

6 Sugiura, A., Mattie, S., Prudent, J. & McBride, H. M. Newly born peroxisomes are a hybrid of mitochondrial and ER-derived pre-peroxisomes. Nature 542, 251–254 (2017). 10.1038/nature21375

7 Hughes, A. L., Hughes, C. E., Henderson, K. A., Yazvenko, N. & Gottschling, D. E. Selective sorting and destruction of mitochondrial membrane proteins in aged yeast. Elife 5 (2016). 10.7554/eLife.13943

8 Schuler, M. H. et al. Mitochondrial-derived compartments facilitate cellular adaptation to amino acid stress. Mol Cell 81, 3786–3802 e3713 (2021). 10.1016/j.molcel.2021.08.021

9 Sugiura, A., McLelland, G. L., Fon, E. A. & McBride, H. M. A new pathway for mitochondrial quality control: mitochondrial-derived vesicles. EMBO J 33, 2142–2156 (2014). 10.15252/embj.201488104

10 Li, X. et al. Mitochondria shed their outer membrane in response to infection-induced stress. Science 375, eabi4343 (2022). 10.1126/science.abi4343

11 Pernas, L. et al. Toxoplasma effector MAF1 mediates recruitment of host mitochondria and impacts the host response. PLoS Biol 12, e1001845 (2014). 10.1371/journal.pbio.1001845

12 Blank, M. L. et al. Toxoplasma gondii association with host mitochondria requires key mitochondrial protein import machinery. Proc Natl Acad Sci U S A 118 (2021). 10.1073/pnas.2013336118

13 Itakura, E., Kishi-Itakura, C. & Mizushima, N. The hairpin-type tail-anchored SNARE syntaxin 17 targets to autophagosomes for fusion with endosomes/lysosomes. Cell 151, 1256–1269 (2012). 10.1016/j.cell.2012.11.001

14 Gutierrez, M. G., Munafo, D. B., Beron, W. & Colombo, M. I. Rab7 is required for the normal progression of the autophagic pathway in mammalian cells. J Cell Sci 117, 2687–2697 (2004). 10.1242/jcs.01114

15 Mindell, J. A. Lysosomal acidification mechanisms. Annu Rev Physiol 74, 69–86 (2012). 10.1146/annurev-physiol-012110-142317

16 Yoshimori, T., Yamamoto, A., Moriyama, Y., Futai, M. & Tashiro, Y. Bafilomycin A1, a specific inhibitor of vacuolar-type H(+)-ATPase, inhibits acidification and protein degradation in lysosomes of cultured cells. J Biol Chem 266, 17707–17712 (1991).

17 Abu-Remaileh, M. et al. Lysosomal metabolomics reveals V-ATPase- and mTOR-dependent regulation of amino acid efflux from lysosomes. Science 358, 807–813 (2017). 10.1126/science.aan6298

18 Sinai, A., Webster, P. & Joiner, K. Association of host cell endoplasmic reticulum and mitochondria with the Toxoplasma gondii parasitophorous vacuole membrane: a high affinity interaction. J Cell Sci 110, 2117–2128 (1997).

19 Chen, W. W., Freinkman, E., Wang, T., Birsoy, K. & Sabatini, D. M. Absolute Quantification of Matrix Metabolites Reveals the Dynamics of Mitochondrial Metabolism. Cell 166, 1324–1337 e1311 (2016). 10.1016/j.cell.2016.07.040

20 Hurley, J. H., Coyne, A. N., Miaczynska, M. & Stenmark, H. The expanding repertoire of ESCRT functions in cell biology and disease. Nature 642, 877–888 (2025). 10.1038/s41586-025-08950-y

21 Cygan, A. M. et al. Proximity-Labeling Reveals Novel Host and Parasite Proteins at the Toxoplasma Parasitophorous Vacuole Membrane. mBio 12, e0026021 (2021). 10.1128/mBio.00260-21

22 Hurley, J. H. ESCRT complexes and the biogenesis of multivesicular bodies. Curr Opin Cell Biol 20, 4–11 (2008). 10.1016/j.ceb.2007.12.002

23 Kimura, S., Noda, T. & Yoshimori, T. Dissection of the autophagosome maturation process by a novel reporter protein, tandem fluorescent-tagged LC3. Autophagy 3, 452–460 (2007). 10.4161/auto.4451

24 Delgado, J. M., Shepard, L. W., Lamson, S. W., Liu, S. L. & Shoemaker, C. J. The ER membrane protein complex restricts mitophagy by controlling BNIP3 turnover. EMBO J 43, 32–60 (2024). 10.1038/s44318-023-00006-z

25 Vietri, M., Radulovic, M. & Stenmark, H. The many functions of ESCRTs. Nat Rev Mol Cell Biol 21, 25–42 (2020). 10.1038/s41580-019-0177-4

26 Li, X. et al. Mitochondria shed their outer membrane in response to infection-induced stress. Science 375, eabi4343 (2022). 10.1126/science.abi4343 PMID - 35025629

27 Rivera-Cuevas, Y. et al. Toxoplasma gondii exploits the host ESCRT machinery for parasite uptake of host cytosolic proteins. PLoS Pathog 17, e1010138 (2021). 10.1371/journal.ppat.1010138

28 Fischer, H. G., Stachelhaus, S., Sahm, M., Meyer, H. E. & Reichmann, G. GRA7, an excretory 29 kDa Toxoplasma gondii dense granule antigen released by infected host cells. Mol Biochem Parasitol 91, 251–262 (1998).

29 Tomita, T. et al. Toxoplasma gondii Matrix Antigen 1 Is a Secreted Immunomodulatory Effector. mBio 12 (2021). 10.1128/mBio.00603-21

30 Coppens, I. et al. Toxoplasma gondii sequesters lysosomes from mammalian hosts in the vacuolar space. Cell 125, 261–274 (2006).

31 Prashar, A. et al. Lysosomes drive the piecemeal removal of mitochondrial inner membrane. Nature 632, 1110–1117 (2024). 10.1038/s41586-024-07835-w

32 Pernas, L., Bean, C., Boothroyd, J. C. & Scorrano, L. Mitochondria Restrict Growth of the Intracellular Parasite Toxoplasma gondii by Limiting Its Uptake of Fatty Acids. Cell Metab 27, 886–897 e884 (2018). 10.1016/j.cmet.2018.02.018

33 Chen, W. W., Freinkman, E. & Sabatini, D. M. Rapid immunopurification of mitochondria for metabolite profiling and absolute quantification of matrix metabolites. Nat Protoc 12, 2215–2231 (2017). 10.1038/nprot.2017.104

34 Terkawi, M. A., Kameyama, K., Rasul, N. H., Xuan, X. & Nishikawa, Y. Development of an immunochromatographic assay based on dense granule protein 7 for serological detection of Toxoplasma gondii infection. Clin Vaccine Immunol 20, 596–601 (2013). 10.1128/CVI.00747-12

35 Li, X., Franz, T., Atanassov, I. & Colby, T. Step-by-Step Sample Preparation of Proteins for Mass Spectrometric Analysis. Methods Mol Biol 2261, 13–23 (2021). 10.1007/978-1-0716-1186-9_2

36 Cox, J. & Mann, M. MaxQuant enables high peptide identification rates, individualized p.p.b.-range mass accuracies and proteome-wide protein quantification. Nat Biotechnol 26, 1367–1372 (2008). 10.1038/nbt.1511

37 Ritchie, M. E. et al. limma powers differential expression analyses for RNA-sequencing and microarray studies. Nucleic acids research 43, e47 (2015). 10.1093/nar/gkv007

38 R: A Language and Environment for Statistical Computing (R Foundation for Statistical Computing, 2017).

